# Hypermorphic TaARF4 shapes wheat architecture by repressing SPL-mediated developmental timing

**DOI:** 10.64898/2026.04.23.720357

**Authors:** Zhenru Guo, Yang Li, Kenan Tan, Twan Rutten, Abdullah Shalmani, Qing Chen, Qingcheng Li, Maolin Peng, Lu Lei, Jie Tang, Yudelsy Antonia Tandron Moya, Markus Kuhlmann, Shuangshuang Zhao, Yongyu Huang, Stefan Ortleb, Ricardo Fabiano Hettwer Giehl, Nicolaus von Wiren, Jochen Kumlehn, Youliang Zheng, Yuming Wei, Ke Wang, Pengfei Qi, Thorsten Schnurbusch

## Abstract

Integrating spatial morphogens with temporal developmental clocks is fundamental to optimizing plant architecture and crop yield, yet their molecular interface remains elusive. Here, we characterize *Branched shoot 1* (*Bsh1*), a semi-dominant wheat mutant exhibiting non-canonical upper-aerial branching and aberrant spike development, caused by a T265I substitution in the DNA-binding domain of the auxin response factor TaARF4-A2. Unlike flanking mutations governing protein stability in basal land plants, this central substitution uniquely converts TaARF4-A2 into an auxin-insensitive hypermorphic repressor. The mutant protein disrupts auxin-cytokinin homeostasis and markedly enhances the native repression of SQUAMOSA PROMOTER BINDING PROTEIN-LIKE (SPL) transcription factors, the core timers of developmental phase transitions. This spatial and temporal uncoupling unleashes upper axillary buds from dormancy, promoting aerial branching and diverting resources from reproductive development. Our findings demonstrate how a conserved spatial morphogen effector was evolutionarily rewired in polyploid wheat to dictate the temporal SPL clock, orchestrating species-specific shoot architectural innovation.

## Introduction

Plant architecture is constructed from repeating units called phytomers, each consisting of a node, an internode, one or more axillary meristems (AxMs), and a leaf. A central governing principle of this organization is apical dominance, where the primary shoot apex suppresses the outgrowth of lateral AxMs to prioritize main stem elongation. The developmental program of phytomers is further shaped by phase transitions that coordinate vegetative and reproductive growth^1,2^. During the vegetative phase, phytomers actively promote leaf growth while suppressing AxM outgrowth to maintain apical dominance. Conversely, upon the transition to reproduction, this pattern is inverted: the inflorescence meristem suppresses leaf (bract) development while promoting AxM to differentiate into lateral branches or spikelets, the reproductive structures that bear florets and grains. In common wheat (*Triticum aestivum* L.; AABBDD, 2n = 6x = 42), this hierarchical control is a fundamental prerequisite for grain yield potential and stability. Basal vegetative nodes produce tillers, productive branches initiated at ground level that can form their own spikes, whereas axillary buds at upper culm nodes are strictly suppressed to minimize resource competition. Deviations from this developmental program lead to aberrant plant architecture^3,4^; however, the molecular mechanisms that maintain dormancy of axillary buds at upper culm nodes remain poorly understood.

Plant development relies on the precise coordination of spatial cues and temporal signals. Unraveling the molecular basis of these developmental programs requires dissecting the interplay of phytohormones and developmental timing pathways. Classically, apical dominance is maintained by the antagonism between basipetal auxin transport, which inhibits AxM outgrowth, and cytokinin (CK), which promotes it^5–8^. Beyond dictating the spatial positioning and patterning of organ formation, auxin also influences developmental rhythms and the timing of organ initiation through dynamic oscillations and local maxima^9–11^. Concurrently, temporal phase transitions (juvenile-to-adult and vegetative-to-reproductive) are primarily governed by the *miR156*–SQUAMOSA PROMOTER BINDING PROTEIN-LIKE (SPL) module. The age-dependent decline of *miR156* allows the progressive accumulation of its target SPL transcription factors, thereby conferring developmental competence to meristems and reshaping plant architecture^12–17^. In rice, for instance, the SPL family member *IPA1* (*OsSPL14*) acts as a master regulator of the "ideal plant architecture" by optimizing tiller number and panicle branching^18,19^. Crucially, although the *miR156*-SPL module has been shown to modulate auxin biosynthesis, transport, and sensitivity^13,20–22^, whether spatial auxin signals and their downstream nuclear auxin pathway (NAP) components can directly regulate the temporal SPL timer remains elusive. Consequently, in the complex polyploid genome of wheat, it remains unclear how auxin signaling is integrated with these developmental regulators to maintain strict dormancy at upper nodes and establish reproductive phytomer identity.

To orchestrate these precise developmental decisions, land plants evolved the NAP, which comprises TIR1/AFB receptors, Aux/IAA repressors, and auxin response factor (ARF) transcription factors^23^. Phylogenomic analyses suggest that the assembly of this pathway was a key innovation that enable the complex coordinated regulation of phytomer development during plant terrestrialization^24–26^. Although a “minimal” precursor system existed in charophyte algae, the subsequent radiation of angiosperms was accompanied by the extensive expansion and functional diversification of these gene families. This diversification transformed the ancestral pathway into a complex regulatory network capable of controlling AxM fates across the plant body^27–30^. Within this expanded system, ARFs diverged into activators and repressors classed with conserved DNA-binding domains that determine transcriptional outputs^31–33^. However, how conserved residues within these domains have been evolutionarily constrained or repurposed to modulate repressive activity in the polyploid crops remains unclear. Specifically, it is unknow how wheat exploits these conserved signaling nodes to orchestrate its distinct architectural traits, such as the strict suppression of aerial branching.

Here, we report the genetic and functional characterization of *Branched shoot 1* (*Bsh1*), a semi-dominant wheat mutant exhibiting a loss of apical dominance and the transformation of dormant upper AxMs into ectopic aerial branches. We show that *Bsh1* phenotypes arises from a gain-of-function T265I substitution in the auxin response factor TaARF4-A2. This substitution reveals a wheat-specific functional dependency that contrasts with the flanking-site stabilization mechanism observed in bryophytes^34,35^. Mechanistically, we demonstrate that the T265I substitution locks TaARF4-A2 into an auxin-insensitive hyper-repressor that disrupts hormonal homeostasis and directly represses the SPL regulatory module. Our results reveal an evolutionary repurposing of a conserved signaling motif that orchestrates reproductive phytomer development and shoot architecture in a polyploid cereal.

## Results

### Semi-dominant *Bsh1* Deregulates Apical Dominance and Confers Developmental Competency to Upper Nodes

To understand how apical dominance in wheat is developmentally controlled, we isolated a semi-dominant mutant, *Branched shoot 1* (*Bsh1*), from an ethyl methanesulfonate (EMS)-mutagenized population of *T. aestivum* cv. Shumai482 (SM482). Phenotypic characterization in both the original SM482 background and subsequently derived near-isogenic lines (NILs; hereinafter referred to as *Bsh1* and wild-type, WT) revealed profound pleiotropic defects. These included attenuated apical dominance, dwarfism, delayed flowering, and a proliferation of leaves, tillers, and both culm and rachis internodes (Fig. 1a–e; Supplementary Figs. 1 and 2). Strikingly, the mutant displayed distinctive spike anomalies: basal spikelets were rudimentary with glumes and lemmas elongating into leaf-like bract structures, and subtending leaf primordia were incompletely suppressed (Fig. 1d, e; Supplementary Fig. 2d). Furthermore, the basal rachis internodes appeared wavy or kinked, whereas the lowermost internode was excessively elongated (Fig. 1e).

**Fig. 1:**
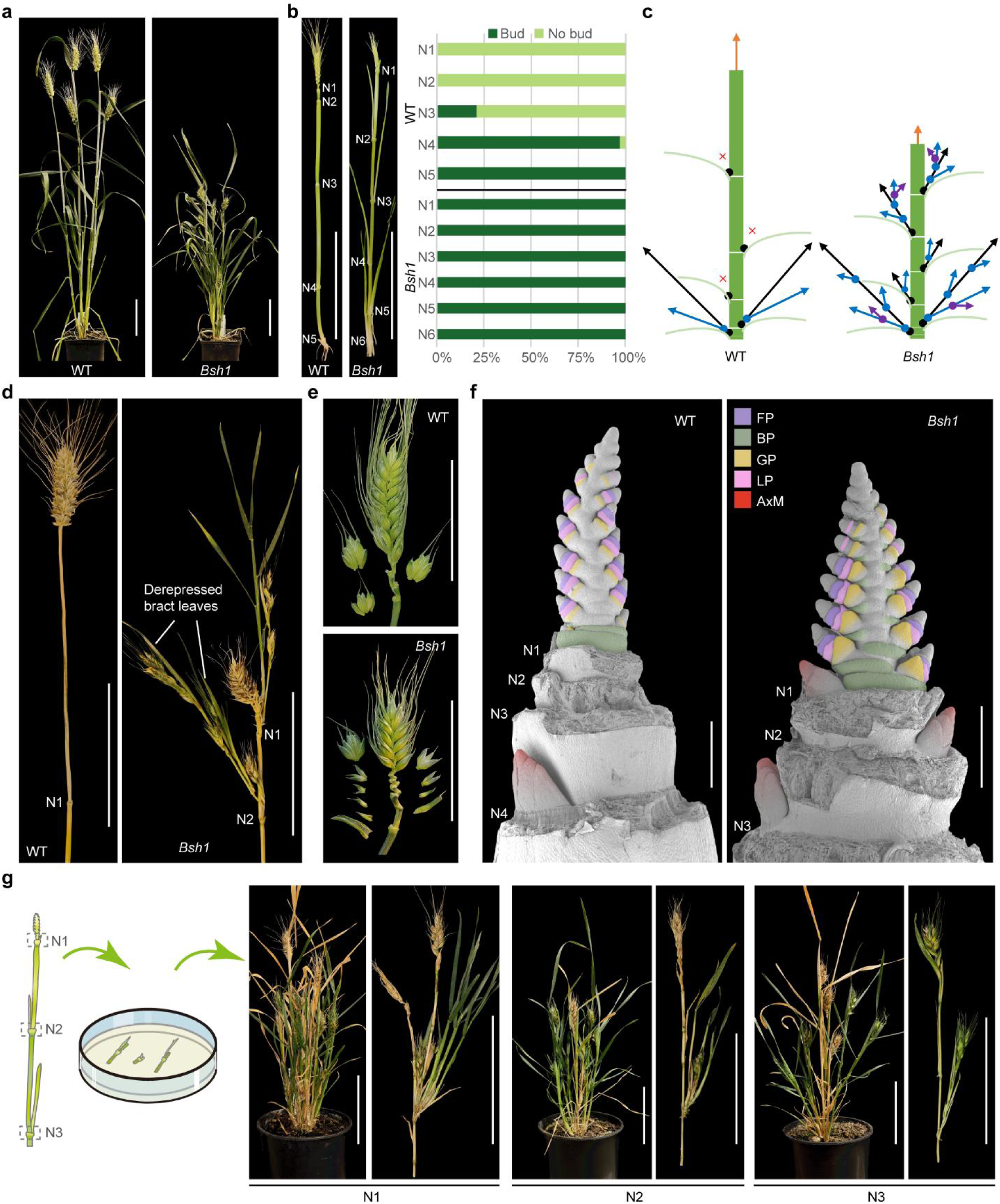
Disrupted apical dominance and phytomer identity in *Bsh1*. **a**, Whole-plant architecture phenotypes of the wild type (WT) and *Bsh1*. Scale bar, 10 cm. **b**, Main culm phenotypes showing nodes N1 to N6 (left) and the proportion of plants developing axillary buds at each node position (right). N1 to N6 denote Node 1 to Node 6. Scale bar, 10 cm. *n* = 33 and 30 biologically independent plants for WT and *Bsh1*, respectively. **c**, Schematic representation comparing the branching architecture and axillary bud outgrowth between WT and *Bsh1*. Black, blue, and purple arrows indicate primary, secondary, and tertiary branches, respectively; red crosses indicate repressed axillary meristems (AxM) in WT. **d**, **e**, *Bsh1* exhibits ectopic aerial branching and derepressed bract leaves at the upper nodes (**d**) and suppressed development of basal spikelets in the spikes (**e**). Scale bar, 5 cm. **f**, Scanning electron microscopy (SEM) analysis of AxM initiation at the floret primordium stage (W3.5). WT shows no visible AxMs at the upper three nodes, whereas *Bsh1* initiates AxMs at every culm node. Scale bar, 500 μm. FP: floret primordium; BP: bract primordium; GP: glume primordium; LP: lemma primordium; AxM, axillary meristem. **g**, Assessment of developmental competency and regeneration potential of the upper three nodes (N1-N3) in *Bsh1*. Schematic of the *in vitro* culm node regeneration assay (left) and representative regenerated plants from the respective nodes (right). Scale bar, 10 cm.

**Fig. 2:**
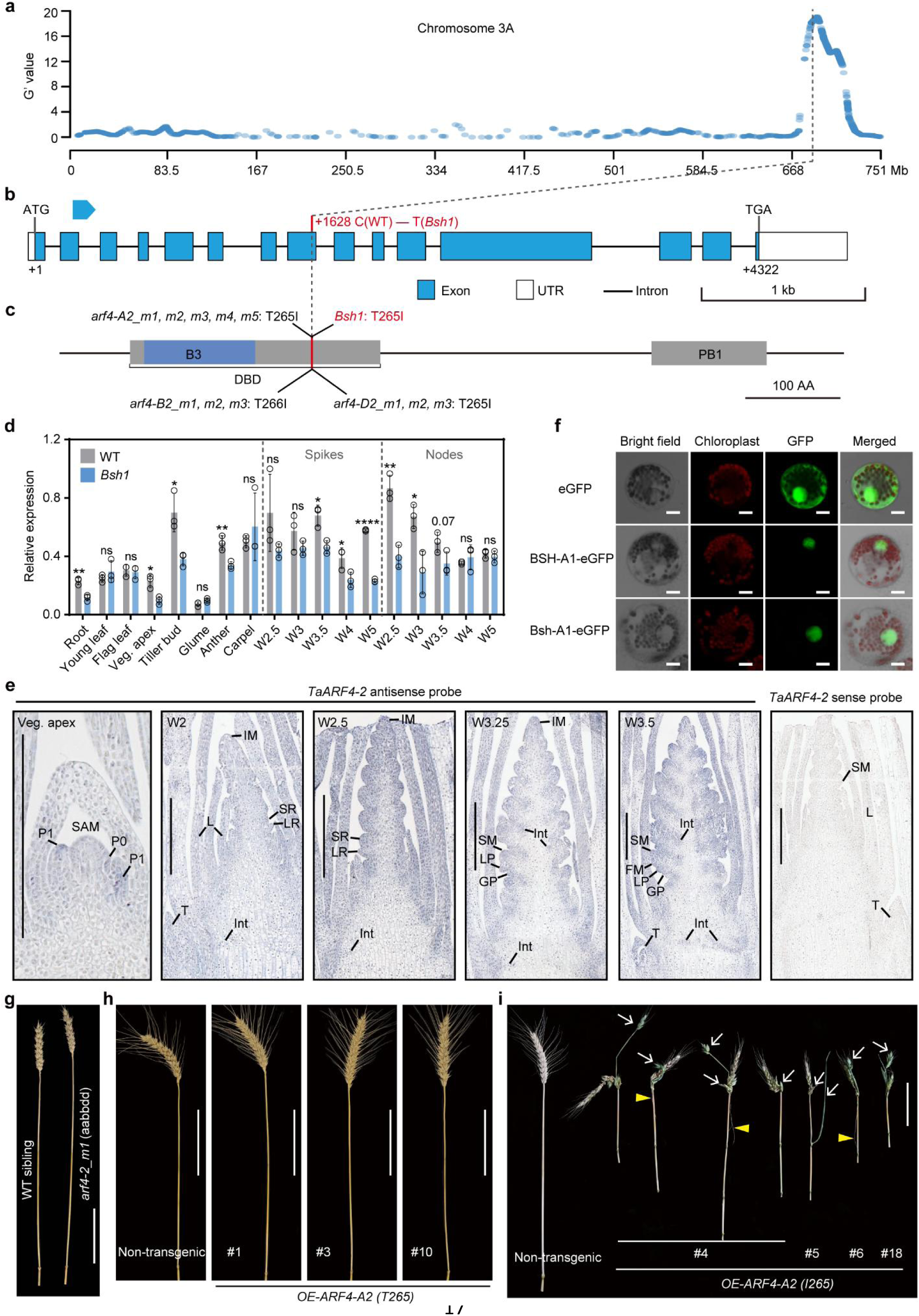
Identification and characterization of *TaARF4-A2* as the causal gene. **a**, Genome-wide G′ statistic plot derived from bulked segregant exome-sequencing. The prominent peak on chromosome 3A indicates the candidate region linked to the mutant phenotype. **b**, Gene structure of *TraesCS3A02G449300* (*TaARF4-A2*). A single nucleotide polymorphism (SNP) is located in exon 8. Exons (blue boxes), 5’ and 3’ UTRs (white boxes), and introns (black lines) are indicated. **c**, Schematic representation of conserved protein domains in *ARF4-2* homoeologs, showing the DNA-binding domain (DBD), B3-type DBD (B3), and PB1 domains. The relative location of the T265I mutations is highlighted in red. **d**, Relative expression levels of *TaARF4-A2* in various tissues analyzed by RT-qPCR. Data are means ± s.d. (*n* = 3 biological replicates; each replicate represents a pool of 5 to 20 individuals). **e**, *In situ* hybridization showing *TaARF4-2* expression patterns at the vegetative apex, early double ridge stage (W2), double ridge stage (W2.5), glume-lemma primordium stage (W3.25), and floret primordium stage (W3.5) using an antisense probe. A sense probe was used as a negative control. SAM, shoot apical meristem; P0/P1, nascent leaf primordia; IM, inflorescence meristem; SR, spikelet ridge; LR, leaf ridge; SM, spikelet meristem; FM, floret meristem; Int, intercalary meristem; L, leaf; T, tiller bud. Scale bar, 500 μm. **f**, Subcellular localization of wild-type and Bsh-A1 proteins fused with eGFP in wheat mesophyll protoplasts. Scale bar, 10 μm. **g**, Phenotypes of the spike and peduncle node in the triple loss-of-function mutant *arf4-2_m1* compared to its WT sibling. Scale bar, 10 cm. **h-i**, Phenotypic characterization of transgenic wheat lines overexpressing the wild-type allele (T265) (**h**) and the *Bsh1* allele (I265) (**i**). White arrows indicate non-canonical aerial branches, yellow arrowheads indicate aerial roots. Scale bar, 10 cm.

Most notably, while the upper three culm nodes (N1–N3) of WT wheat are typically dormant, *Bsh1* initiated functional axillary buds at every node position. These ectopic buds elongated into high-order branches bearing abnormal inflorescences, including aerial roots (Fig. 1b-d; Supplementary Fig. 2b, c). To investigate the ontogeny of these axillary buds, we performed comparative dissection and histological analysis of developing spikes and culm nodes (Fig. 1f and Extended Data Fig. 1). In *Bsh1*, ectopic AxMs were detectable as early as the Waddington 2.5 (W2.5) stage^36^, a developmental window where no meristematic activity is observed in the corresponding WT upper nodes. These meristems subsequently differentiated into axillary buds containing developing spikes; occasionally, buds also emerged at the spike collar. Further examination using longitudinal sections of resin-embedded apices captured the precise initiation dynamics: during the vegetative phase, dome-like meristematic protrusions emerged in the axil of every nascent leaf primordium in *Bsh1*, subsequently progressing into distinct axillary buds. In contrast, the WT showed no signs of meristem initiation at the prospective upper three nodes, even after the transition to spike differentiation (Supplementary Fig. 3a, b). To assess the developmental competency of these ectopic buds, we cultured excised upper nodes (N1–N3) *in vitro*, using the basal rachis node (N0) as a negative control. After three weeks of culture and subsequent transplantation to soil, only *Bsh1* nodes regenerated into complete plants retaining the hyper-branching phenotype (Fig. 1g and Supplementary Fig. 3c-d). These results confirm that the mutant buds function as viable, high-position tillers with autonomous developmental potential.

Collectively, these data demonstrate that *Bsh1* mutation(s) drive the excessive generation of leaf primordia and phytomers, facilitates axillary bud formation and outgrowth at culm nodes, while simultaneously suppressing basal spikelet identity in favor of bract growth in the inflorescence.

### A Hypermorphic Mutation in *TaARF4-A2* Underlies the *Bsh1* Phenotypes

To elucidate the genetic basis of the *Bsh1* phenotypes and effectively minimize the interference of EMS-induced background mutations, we strategically advanced the mapping population derived from the cross between *Bsh1* and QZ212 by continually selecting and selfing heterozygous plants (Supplementary Fig. 1d). Phenotypic assessment of 132 individuals from an advanced segregating family revealed a segregation of 24 plants displaying the homozygous mutant phenotype, 66 exhibiting an intermediate (heterozygous) phenotype, and 42 showing normal architecture. This distribution fits a 1:2:1 ratio (χ^2^ = 4.91, *p* = 0.086), suggesting that the phenotype is governed by a single, semi-dominant nuclear locus.

We initially mapped the *Bsh1* locus to the long arm of chromosome 3A utilizing Bulked Segregant Exome-sequencing (BSE-seq) on extreme phenotypic pools explicitly selected from the segregating F_4_ generation (Fig. 2a and Extended Data Fig. 2a). To fine-map the causal gene, we developed polymorphic molecular markers based on local SNPs and screened a large, high-resolution mapping population comprising 3,009 individuals from the advanced F_2:5_ and F_2:6_ generations. This effort delimited the *Bsh1* locus to a 275-kb physical interval flanked by markers *m10* and *m14*, with the mutant phenotype completely co-segregating with the central marker *m13* (Extended Data Fig. 2b-c and 3a). Crucially, progeny testing of residually heterozygous lines (RHLs) derived from key recombinants independently validated this physical interval, perfectly associating the distinctive *Bsh1* architectural defects with the *m13* genotype (Extended Data Fig. 3b–c). Reference genome annotation revealed six high-confidence protein-coding genes within this interval. Subsequent Sanger sequencing of all candidate genes in the parental lines identified only a single causative polymorphism: A C-to-T transition in the eighth exon of *TraesCS3A02G449300* (Fig. 2b and Extended Data Fig. 2d). This non-synonymous mutation results in a Threonine (T) to Isoleucine (I) substitution at the conserved residue 265 (T265I) (Fig. 2b–c and Extended Data Fig. 4a). Phylogenetic analysis established this gene as the wheat ortholog of the well-characterized auxin response factors *AtARF2*^37^ in *Arabidopsis* and *OsARF4*^38^ in rice; consequently, we assigned the symbol *TaARF4-A2* for the gene name (Extended Data Fig. 2e). Following updated guidelines for gene nomenclature in wheat^39^, we designated the dominant mutant allele as *Bsh-A1* and the wild-type allele as *BSH-A1*.

To understand the developmental context of *TaARF4-A2* action, we analyzed its spatiotemporal expression profile. Quantitative Real-Time PCR (qRT-PCR) revealed that *TaARF4-A2* is constitutively expressed, with high levels in developing young spikes, early culm nodes, and tiller buds (Fig. 2d). mRNA *in situ* hybridization (ISH) further refined this pattern, showing that *TaARF4-2* transcripts accumulated in nascent leaf primordia and the shoot apical meristem (SAM) during the vegetative stage. Upon the reproductive transition, expression persisted in vegetative structures (tillering buds, leaves) and reproductive structures (glume/lemma primordia, inflorescence meristems, spikelet meristems, and floral meristems), as well as in the intercalary meristems of culm and rachis internodes (Fig. 2e). These observations were corroborated by spatial transcriptomic datasets from tetraploid wheat (*T. turgidum* ssp. *durum* cv. Kronos) spikes^40^, in which *TaARF4-2* displayed highly consistent constitutive expression patterns (Supplementary Fig. 4a-b). Similarly, spatial transcriptomic profiling of developing barley spikes demonstrated that the two *HvARF4* paralogs share a highly conserved expression domain with their wheat orthologs^41,42^. This broad spatiotemporal foot print aligns with the pleiotropic defects observed in *Bsh1*. Subcellular localization analysis using eGFP-tagged proteins in wheat protoplasts further confirmed that the T265I mutation did not affect the protein’s subcellular nuclear localization (Fig. 2f).

Guided by these expression data, we sought to delineate the functional nature of the *Bsh1* allele—specifically, whether it represents a loss-of-function (LOF) or gain-of-function mutation. We first characterized a targeted TILLING population for *TaARF4-A2*, identifying four premature termination alleles (putative LOF) and four missense alleles. In addition, we analyzed 13 mutants targeting its homoeologs (*TaARF4*-*B2*: *TraesCS3B02G486000*; *TaARF4-D2: TraesCS3D02G442000*) and closely related paralogs (*TaARF4*-*A1*: *TraesCS3A02G442000*; *TaARF4-B1: TraesCS3B02G475800*), comprising nine premature termination alleles and four missense alleles. None of these mutants exhibited the aerial branching phenotype in upper nodes (Extended Data Figs. 4-5). Consistently, a triple LOF mutant (*arf4-2_m1*) generated by crossing exhibited traits opposite to *Bsh1*, including reduced tillering and increased plant height but no aerial branching of upper nodes (Fig. 2g and Extended Data Fig. 6). Having ruled out LOF as the cause, we next tested whether the T265I substitution is sufficient to confer the *Bsh1* phenotype using transgenic assays. In striking contrast to the LOF mutants, transgenic lines overexpressing the mutant allele (*Bsh-A1*, I265) in *T. aestivum* cv. Fielder recapitulated the *Bsh1* syndrome, including upper aerial branching and spike deformities (Fig. 2i and Extended Data Fig. 7d–f); whereas lines overexpressing the WT allele (T265) remained normal (Fig. 2h and Extended Data Fig. 7a-c). Consistent with this, we identified 11 independent EMS mutants from distinct populations that carried identical T-to-I substitutions in the AA, BB, or DD homoeologs of *TaARF4-2*, all of which exhibited aerial branching in upper nodes and accordingly reproduced the *Bsh1* syndrome (Fig. 2c and Extended Data Fig. 8). Together, these converging lines of evidence demonstrate that the T265I substitution confers a hypermorphic gain-of-function protein. Moreover, heterologous expression of the *Bsh-A1* mutant allele (I265) in *Arabidopsis* significantly impeded the vegetative-to-reproductive transition, as evidenced by delayed bolting and an increased rosette leaf number (Extended Data Fig. 9). While overexpression of the wild-type *TaARF4-A2* (T265) resulted in only a marginal, non-significant delay in flowering compared to the Col-0 background, the *Bsh-A1* variant acted as a potent repressor, profoundly extending the vegetative phase. These results demonstrate that the hypermorphic potency of the T265I substitution is functionally conserved across divergent plant lineages.

### T265I Confers Site-specific Hyper-repression and Dominant Auxin Insensitivity

To elucidate the molecular basis of the semi-dominant *Bsh1* phenotypes, we first examined the structural consequences of the T265I substitution. Structural modeling using AlphaFold3 predicted that the mutation does not appreciably alter the overall architecture of the DNA-binding domain (DBD). Instead, it replaces a polar threonine with a bulky hydrophobic isoleucine within a solvent-exposed and evolutionarily conserved loop (Fig. 3a and Supplementary Fig. 5a).

**Fig. 3:**
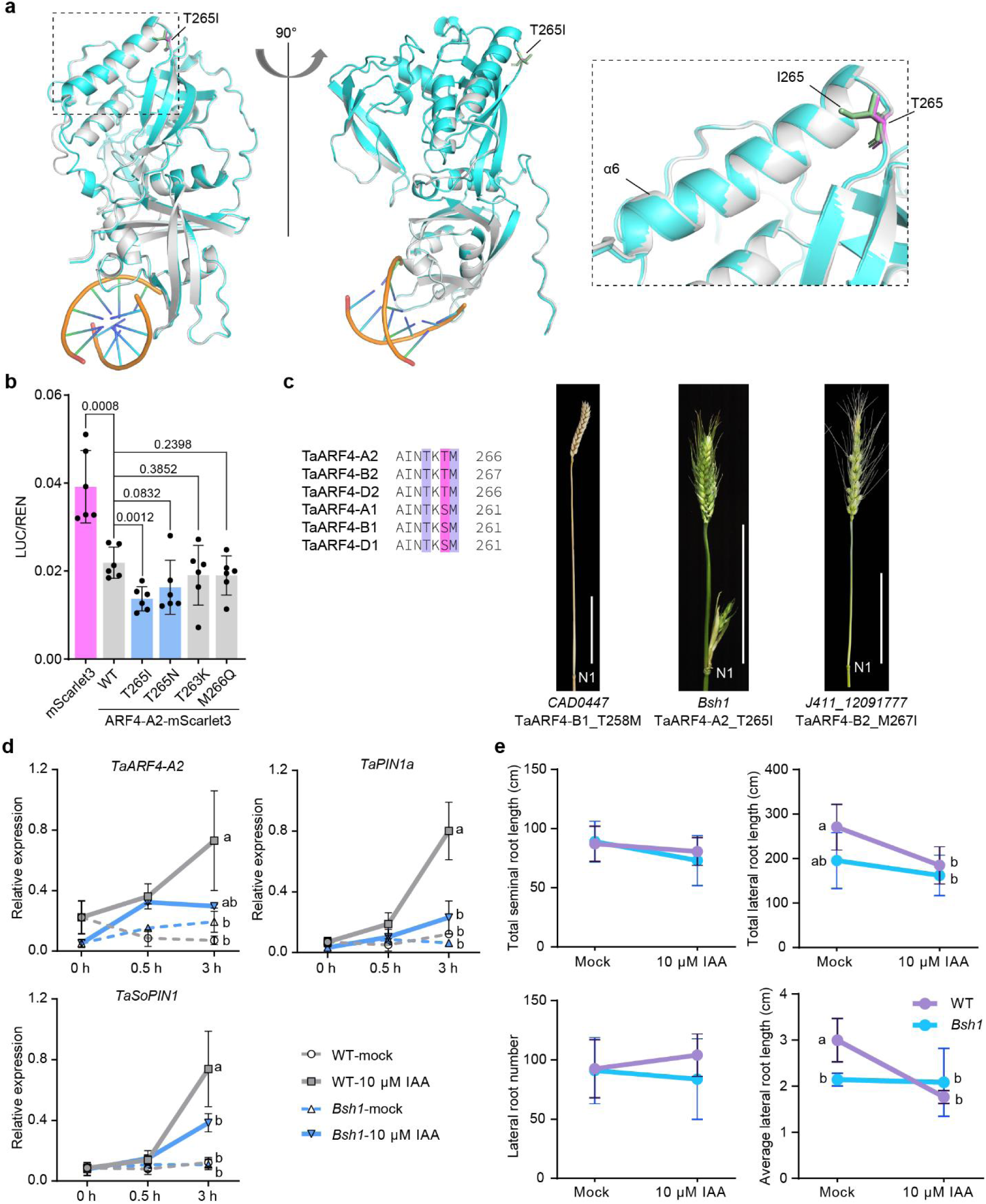
T265I substitution confers dominant auxin insensitivity and hyper-repressive activity. **a**, Structural superimposition of the predicted DNA-binding domains (DBD) of TaARF4-A2 WT (grey) and the Bsh-A1 (I265) variant (cyan), generated using AlphaFold3. Residue T265 (WT) and I265 (mutant) are shown as sticks. The enlarged view highlights the solvent-exposed loop harboring residue 265, indicating that the T265I substitution does not cause detectably alterations in local secondary structure or overall DBD folding. **b**, Dual-luciferase reporter assays assessing the functional specificity of T265 relative to adjacent conserved residues. TaARF4-2 variants were co-expressed with a *TaSPL-D14* promoter-driven LUC reporter. T265I and T265N substitutions confer enhanced repression activity, whereas mutations in neighboring residues (T263K and M266Q) do not. Data are means ± s.d. (*n* = 6 biologically independent samples). *P* values were determined by two-sided Student’s *t t*est. **c**, Left, sequence alignment of six wheat TaARF4-2 homoeologs (A1/B1/D1 and A2/B2/D2) showing complete conservation of the DBD loop region. The Bsh1 mutation site (T265, purple) and adjacent residues targeted in TILLING control lines (T258 and M267, blue) are indicated. Right, representative spike and peduncle node phenotypes demonstrating that TILLING mutants of adjacent conserved residues retain normal architecture, in contrast to the basal spikelet deformities and nodal branching observed in *Bsh1*. Whole-plant phenotypes are shown in Supplementary Fig. 8. Scale bars, 10 cm. **d**, Relative transcript levels of *TaARF4-A2*, *TaPIN1a*, and *TaSoPIN1* in roots from hydroponically grown seedlings measured by RT-qPCR (*n* = 3 biologically independent samples). Auxin-induced activation of *PIN* genes observed in WT is abolished in *Bsh1*. Different letters indicate statistically significant differences (one-way ANOVA with Tukey’s multiple comparisons test, *p* < 0.05). Data are means ± s.d. **e**, Root growth response of WT and *Bsh1* seedlings grown for 11 days in hydroponics culture supplemented with mock solution or 10 μM indole-3-acetic acid (IAA). Quantification shows reduced sensitivity of *Bsh1* lateral root elongation to high auxin concentrations (*n* ≥ 5 biologically independent plants). Data are means ± s.d.

Recent studies in basal land plants, including *Physcomitrium patens* and *Marchantia polymorpha*, have identified this loop as a regulatory hotspot influencing ARF protein stability, with substitutions at flanking residues markedly enhancing protein accumulation^34,35^. To test whether analogous substitutions affect TaARF4-A2 activity in wheat, we performed dual-luciferase assays using a *TaSPL-D14* promoter-driven reporter (validated as a direct target in Fig. 5). Substitutions mimicking the basal-plant stabilizing mutations at adjacent residues (T263K and M266Q) did not alter transcriptional repression relative to the wild-type TaARF4-A2 effector (Fig. 3b). Instead, enhanced repression was strictly dependent on the central T265 residue: both the T265I substitution and a T265N mutation (mimicking the maize *Truffula* gain-of-function allele, S281N) functioned as profound hyper-repressive variants (Fig. 3b).

These results indicate strong positional specificity at residue T265 in wheat TaARF4. To assess whether this specificity extends *in vivo*, we analyzed TILLING lines carrying substitutions at conserved neighboring residues, including *TaARF4-B1* (T258M) and *TaARF4-B2* (M267I). Although the DBD loop is fully conserved among wheat homoeologs (Fig. 3c, left; Extended Data Fig. 4a), these adjacent substitutions neither altered the predicted DBD structure (Supplementary Fig. 5b-c) nor affected spike or peduncle node architecture, which remained indistinguishable from the WT (Fig. 3c, right; Extended Data Fig. 5a,c).

Having established Bsh-A1 (i.e. TaARF4-A2_I265_) as a site-specific hyper-repressor, we next examined its physiological impact on auxin signaling. Because ARFs regulate auxin transport and feedback networks, we analyzed the expression of canonical auxin efflux carriers. RT-qPCR of hydroponically grown wild-type roots showed that exogenous indole-3-acetic acid (IAA) robustly induced *TaARF4-A2*, *TaPIN1a*, and *Sister of PIN1* (*TaSoPIN1*) expression. In contrast, this auxin-responsive transcriptional activation was abolished in *Bsh1* (Fig. 3d).

This profound transcriptional blockade of the auxin transport machinery suggested that *Bsh1* is fundamentally “deaf” to auxin signaling. To determine whether this transcriptional attenuation translates into physiological auxin insensitivity, we assessed root growth responses to exogenous IAA. Primary seminal root elongation on agar plates showed only minor differences between genotypes (Supplementary Fig. 6a), suggesting no general growth defect. However, lateral root elongation under hydroponic conditions was strongly inhibited by high IAA concentrations in the WT, whereas *Bsh1* lateral roots exhibited markedly reduced sensitivity (Fig. 3e and Supplementary Fig. 6b). Finally, to confirm that this hormone insensitivity was systemic and inextricably linked to the architectural defects, we attempted to chemically rescue the mutant phenotype. Soil-drenching treatments with exogenous IAA or the auxin transport inhibitor *N*-1-naphthylphthalamic acid (NPA) completely failed to suppress the ectopic aerial branching in *Bsh1* (Supplementary Fig. 6c-e). Collectively these molecular and physiological analyses demonstrate that the T265I substitution locks TaARF4-A2 into a dominant, auxin-insensitive hyper-repressor that systematically disrupts endogenous phytohormonal feedback.

### Transcriptomic Reprogramming and Hormonal Imbalance Driven by the Bsh1-A1 Hypermorphic Repressor

To elucidate how the Bsh-A1 variant orchestrates spike deformity and ectopic axillary bud formation transcriptionally, we performed RNA sequencing (RNA-seq) on young spikes and upper culm nodes (N1–N3) across three critical developmental stages: the double-ridge (DR/W2.5), glume primordium (GP/W3), and floret primordium (FP/W3.5) stages (Supplementary Fig. 7a). Principal component analysis (PCA) showed that PC1 and PC2 accounted for 67.09% of the total variance, distinguishing tissue types and highlighting significant genotypic divergence (Supplementary Fig. 7b). We identified 14,820 differentially expressed genes (DEGs), with the most pronounced transcriptional upheaval occurring at the W2.5 stage in nodes (5,033 upregulated; 1,479 downregulated) and spikes (4,279 upregulated; 2,000 downregulated) (Supplementary Fig. 7c). This timing coincides precisely with the histological emergence of ectopic meristematic domes (see Extended Data Fig. 1), indicating that the Bsh1-A1 variant triggers widespread transcriptional reprogramming at the very onset of AxM initiation. Venn diagram analysis identified a core set of 5,301 DEGs shared between tissues, reflecting the sustained and systemic influence of the mutation (Supplementary Fig. 7d). To dissect regulatory patterns, we categorized these DEGs into ten expression clusters (Fig. 4a, Supplementary Fig. 7e). Gene Ontology (GO) enrichment analysis revealed a striking overrepresentation of auxin-related biological processes across multiple clusters (Fig. 4b), pointing to a pervasive collapse of auxin homeostasis.

**Fig. 4:**
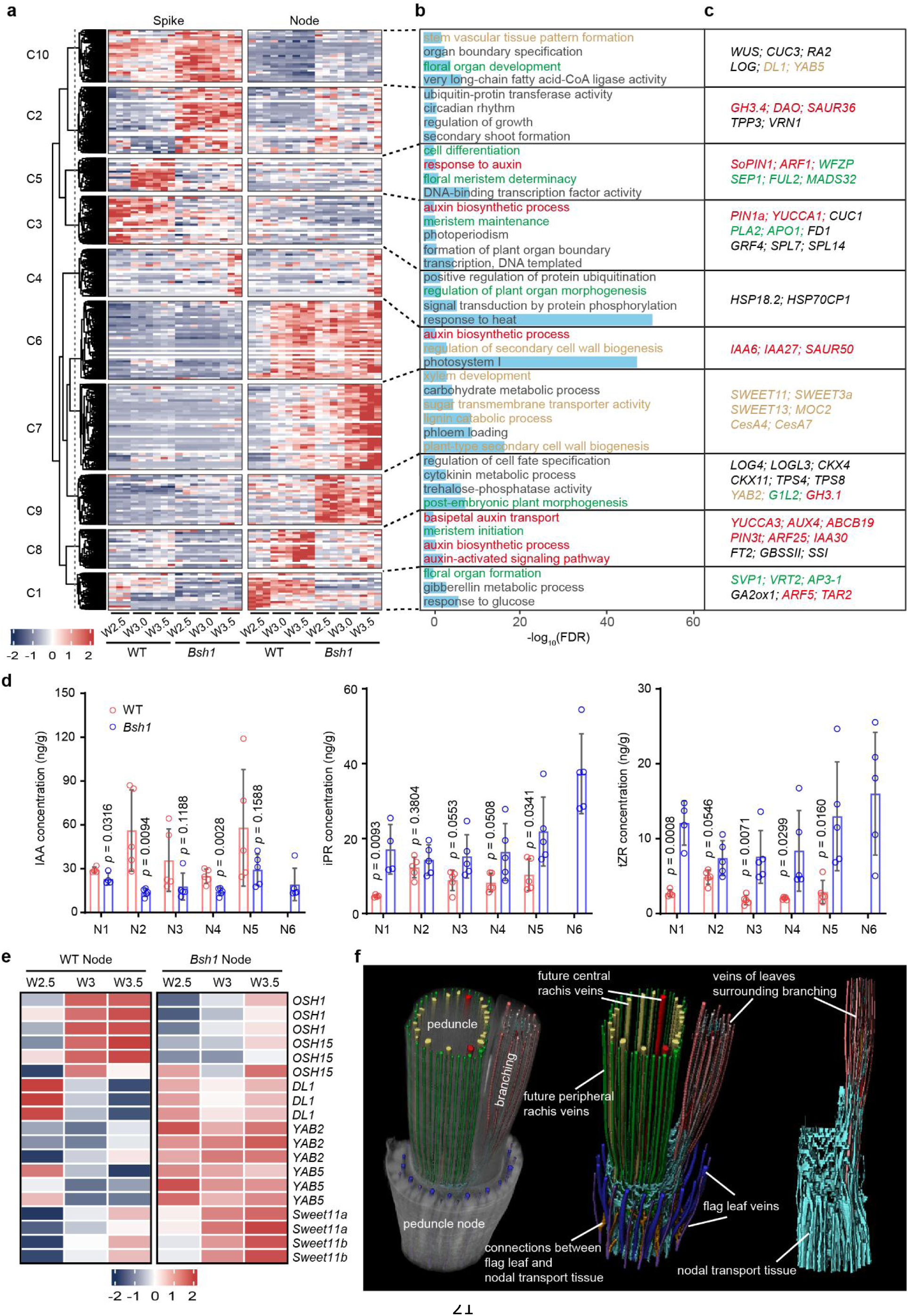
Transcriptional reprogramming and physiological alterations in *Bsh1*. **a**, Clusters of co-expressed genes in spikes and nodes across three developmental stages. **b-c**, Selected Gene Ontology (GO) enrichment terms and representative wheat orthologs from the identified clusters. Genes related to auxin signaling, floral organ development, and vasculature are highlighted in red, green, and gold, respectively. **d**, Concentrations of endogenous auxin and cytokinin in different culm nodes at approximately stage W8. IAA, indole-3-acetic acid; iPR: isopentenyladenosine; tZR, *trans*-zeatin riboside. Data are means ± s.d. (*n* = 4 or 5 biological replicates; each replicate represents a pool of 5 individuals). *P* values were determined by two-sided Student’s *t t*est. **e**, Heatmap showing the relative expression (scaled TPM) of representative vasculature-related genes. **f**, Three-dimensional (3D) reconstruction of vascular bundles connecting the axillary bud at peduncle node of *Bsh1*.

Consistent with the T265I mutation functioning as a hypermorphic “lock” on the ARF repressor, this transcriptomic disruption was characterized by a systemic suppression of the auxin-promoting machinery, coupled with a shifted cytokinin ratio. Key auxin biosynthesis genes, including the rate-limiting *TaYUCCA1/3*^43^ and upstream *TaTAA1/TAR2*^44,45^, were downregulated. Furthermore, the expression of major polar auxin transport carriers—*TaPIN1a*, *TaSoPIN1*, *TaABCB14*, and *TaAUX4*^46–51^—was significantly suppressed (Fig. 4c, Supplementary Fig. 8a). Concomitantly, genes involved in auxin degradation and conjugation were markedly upregulated, including *DIOXYGENASE FOR AUXIN OXIDATION* (*DAO*)^52^ and multiple Group II *GH3* family members (Ta*GH3.1*, *GH3.4*, *GH3.11*)^53,54^ (Fig. 4c, Supplementary Fig. 8a). This coordinated enhancement of catabolism and repression of synthesis/transport creates a “low-auxin” environment, which we subsequently confirmed via direct hormonal quantification (Fig. 4d). The dysregulation of downstream signaling components (e.g., *SAUR32*, *IAA13*, *ARF5*) further reflects a widespread disruption of auxin signaling. Mirroring the detected decrease in auxin, the release of nodal dormancy was driven by altered cytokinin dynamics^7,8^. In *Bsh1* nodes, we observed the significant upregulation of cytokinin activation genes, including *LONELY GUY* (*TaLOG, LOGL3, LOGL6*)^55^ (Supplementary Fig. 8b). While LOG enzymes typically catalyze the conversion of ribosylated precursors into active free forms, hormonal quantification revealed elevated levels of the ribosylated precursors *trans*-zeatin riboside (tZR) and isopentenyladenosine (iPR) in mutant nodes (Fig. 4d). This concurrent accumulation of both signaling components and biosynthetic transcripts—including the compensatory induction of *TaCKX* degrading enzymes—likely reflects a pervasive, feedback-driven activation of the entire cytokinin metabolic network, potentially triggered by the disrupted auxin signaling landscape.

This profound hormonal shift ultimately translates into structural remodeling, the suppression of reproductive identity, and the circumvention of canonical branching pathways. In the mutant nodes, we observed an inversion of the module governing meristem maintenance: members of the indeterminacy-maintaining Class I KNOX subclade (*OSH1*, *OSH15*)^56,57^ were downregulated, while lateral organ-promoting *YABBY* genes (*DL1*, *YAB2*, *YAB5*)^58^ were upregulated (Fig. 4e). To support the metabolic demands of this ectopic outgrowth, sugar transporters (*SWEET11a/b*) and vascular patterning genes were enriched, providing the molecular basis for the peripheral vascular bundles that functionally connect aerial branches to the main nodal vasculature (Fig. 4f and Supplementary Videos 1-2). Concurrently, the *Bsh1* spike phenotype is underpinned by the severe repression of key reproductive regulators. The hypermorphic allele was associated with the downregulation of floral identity genes (Supplementary Fig. 8c), including MADS-box members (the *SEP* family, *VRT2*, *SVP1*, *FUL2*), the spikelet identity and determinacy gene *FZP*, and growth regulators (*TaGRF3/4*). The delayed heading phenotype also correlates with the downregulation of *PLA2* (plastochron regulation) and flowering-time integrators (*TaFD1*, *TaFT2/HD3a*). Finally, an interrogation of the canonical branching pathway revealed a striking paradox: homologs of classic branching inhibitors, such as *TB1*^59^, *GT1*^60^, and *Tru1*^61^ were upregulated in *Bsh1*, while the branching promoter *MOC1*^62^ was downregulated (Supplementary Fig. 8d). This expression pattern—which would typically dictate a “no-branching” phenotype—is pivotal, as it demonstrates that the TaARF4-2-mediated aerial upper-branching pathway operates independently of the classical *MOC1*/*TB1* framework.

### TaARF4-A2 Directly Hyper-represses the SPL Module to Rewire Developmental Architecture

To identify the downstream effectors underlying the architectural reprogramming observed in *Bsh1*, we examined our comparative transcriptome data. Multiple members of the SPL family were downregulated, with homoeologs of *TaSPL7* and *TaSPL14* showing pronounced repression in both spikes and nodes of *Bsh1* (Fig. 5a and Extended Data Fig. 10a). While a concomitant upregulation of *miR156* in the *Bsh1* shoot apex (Fig. 5b) likely contributes to this broad repression of SPL module, we next investigated whether the hypermorphic TaARF4-A2 repressor might also directly target and regulate these *SPL* genes.

**Fig. 5:**
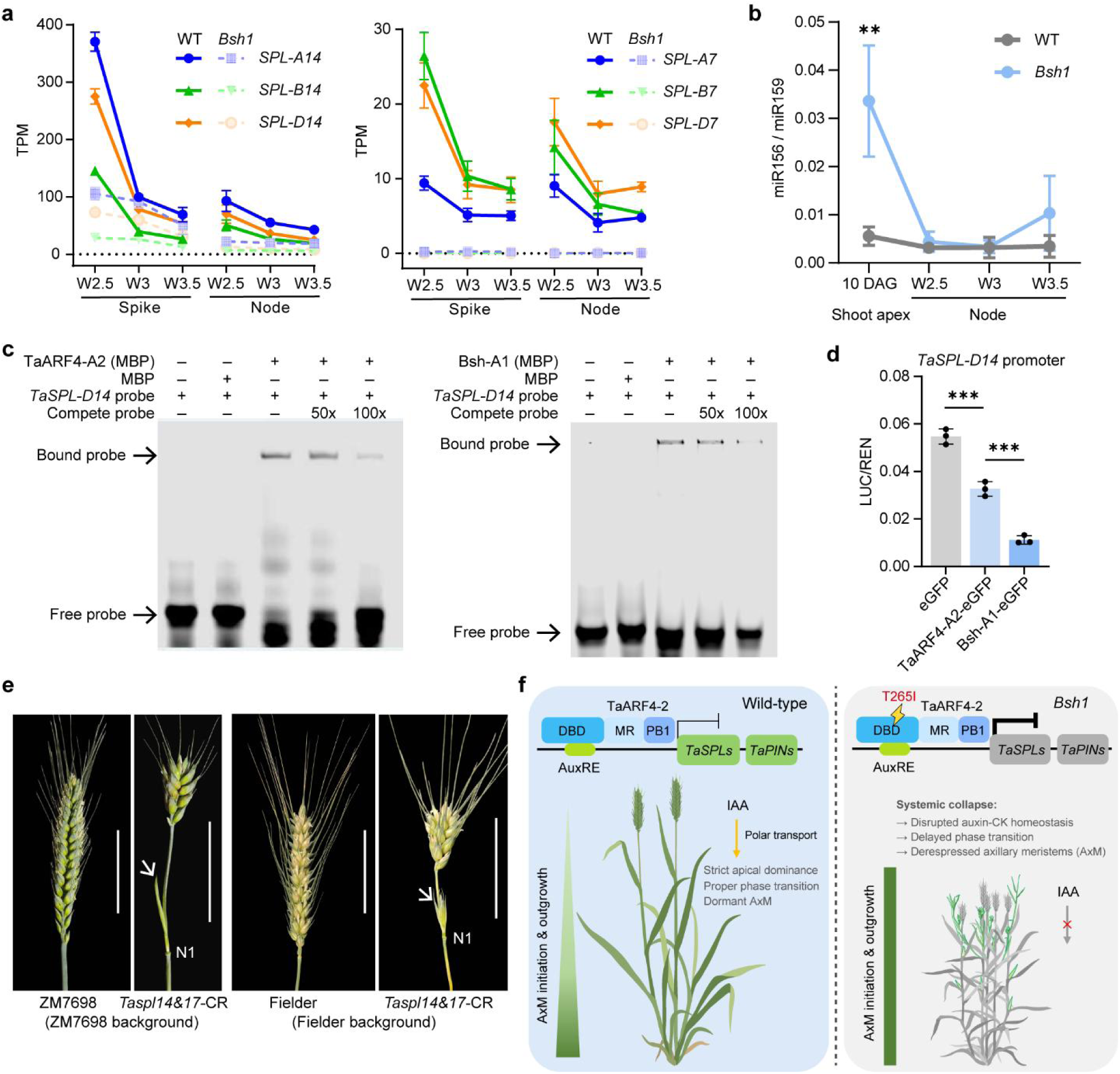
TaARF4-A2 directly binds and hyper-represses *SPL* targets to rewire developmental architecture. **a**, RNA-seq expression profiles (transcripts per million, TPM) of *TaSPL14* (left) and *TaSPL7* (right) homoeologs in developing spikes (W2.5, W3, W3.5) and nodes of WT and *Bsh1*, showing a profound transcriptional shutdown of these key regulators in the mutant. **b**, Stem-loop RT-qPCR analysis of mature miR156 levels relative to miR159 (internal control) in the shoot apex and nodes, demonstrating the cascaded disruption of the developmental timing network. Data are means ± s.d. (*n* = 3 biological replicates). **c**, Electrophoretic mobility shift assay (EMSA) demonstrating that both wild-type MBP-TaARF4-A2 and *Bsh1* mutant (MBP-Bsh1-A1) proteins directly recognize and bind to the canonical auxin response element within the target *TaSPL-D14* promoter *in vitro*. Unlabeled competitor probes (50× and 100×) were added to confirm specific binding. (+) and (−) indicate the presence or absence of components. Black arrows indicate the bound protein-DNA complexes (shifted bands) and free probes. **d**, Dual-luciferase reporter assays showing functional target repression using the *TaSPL-D14* promoter. Compared to the moderate repression by the wild-type TaARF4-A2-eGFP, the T265I mutation in Bsh-A1-eGFP exerts an extreme hyper-repressive effect *in vivo*. Data are means ± s.d. (*n* = 3 biologically independent samples). \*\*\**P* < 0.001, determined by two-sided Student’s *t*-*t*est. **e**, Genetic validation using *Taspl14&17* CRISPR knockout sextuple mutants (*Taspl14&17-CR*) in two independent genetic backgrounds (ZM7698 and Fielder). To overcome functional redundancy, the loss of the entire *TaSPL14/17* module results in abnormal spike architecture and ectopic axillary bud at the peduncle node (white arrows), effectively phenocopying the architectural defects of *Bsh1*. Scale bars, 5 cm. **f**, proposed working model for the species-specific rewiring of auxin signaling by the TaARF4-2-SPL module. **Left:** In the wild-type, TaARF4-2 functions as a moderate repressor of *TaSPLs* and *TaPINs* to maintain canonical apical dominance, proper phase transitions, and polar auxin transport. **Right:** The single-amino-acid T265I substitution transforms TaARF4-2 into a constitutive hyper-repressor (indicated by the bold inhibition line). This “locked” repressor simultaneously shuts down the SPL-mediated phase transition timer and collapses the broader auxin-cytokinin homeostasis networks, ultimately unleashing the ectopic outgrowth of latent axillary meristems.

A fundamental prerequisite for such direct transcriptional regulation is spatiotemporal co-expression. Indeed, spatial transcriptomic profiling of developing spikes revealed that the expression domains of *TaSPL7* and *TaSPL14* extensively overlap with those of *TaARF4* (Supplementary Fig. 4c, d), establishing the requisite biological context for their *in vivo* interaction. Guided by published ATAC-seq data^63^ showing high chromatin accessibility in the proximal promoter regions of *TaSPL7* and *TaSPL14*, we identified canonical ARF-binding motifs within these open chromatin regions (Extended Data Fig. 10b-c). Electrophoretic mobility shift assays (EMSA) confirmed that both the wild-type and the mutant Bsh-A1 (TaARF4-A2_I265_) proteins retain the intrinsic physical capacity to directly recognize and bind to these specific motifs *in vitro* (Fig. 5c and Extended Data Fig. 10d).

To evaluate the functional consequence of this direct physical binding, we performed dual-luciferase reporter assays using the *TaSPL-D14* promoter. Strikingly, while the wild-type TaARF4-A2_T265_ exerted a moderate repressive effect, the Bsh-A1 protein acted as a drastic hyper-repressor, virtually extinguishing reporter expression (Fig. 5d). These molecular data establish *TaSPL14* and *TaSPL7* as direct downstream targets of TaARF4-A2, confirming that the T265I substitution confers a hyper-repressive efficacy without abolishing DNA-binding capability. Rather than disrupting promoter recognition, the T265I mutation appears to trap the DNA-bound TaARF4-A2 in a highly stable, hyper-repressive conformation, preventing the dynamic release or turnover typically required for proper transcriptional regulation.

To genetically test whether repression of the SPL module contributes to the *Bsh1* phenotype, we faced the inherent challenge of extensive functional redundancy in the hexaploid wheat genome. *SPL14* and its closest paralog, *SPL17*, exhibit partially overlapping functions, and single *spl14* mutants often display mild or masked phenotypes due to compensation by *SPL17*^16,64^. To circumvent this redundancy, we evaluated previously reported *spl14 spl17* CRISPR-Cas9 sextuple knockout lines (*Taspl14&17-CR*)^63,65,66^.

Targeted disruption of the entire TaSPL14/17 module in two independent genetic backgrounds resulted in ectopic axillary bud or spikelet-like structures at the peduncle node (N1), accompanied by altered spike morphology and increased internode and tiller numbers (Fig. 5e and Extended Data Fig. 10e, f), partially phenocopying *Bsh1*. While *Bsh1* exhibits a more severe, systemic syndrome due to the simultaneous hyper-repression of multiple SPL clades (e.g., *TaSPL7*) and broader auxin network collapse, these epistasis-like genetic data provide compelling evidence that the TaARF4-2–SPL regulatory axis constitutes a primary molecular checkpoint governing apical dominance and branching architecture in wheat. Together, these findings support a comprehensive working model (Fig. 5f): the species-specific recruitment of the TaSPL module by TaARF4-2 maintains canonical developmental phase transitions, whereas the T265I hyper-repressor mutation short-circuits this network, unleashing latent AxMs thereby profoundly rewiring the plant architecture.

## Discussion

The strict maintenance of apical dominance is a hallmark of cereal architecture, optimizing resource allocation by suppressing upper axillary buds. Our characterization of *Bsh1* uncovers the molecular linchpin enforcing this constraint, revealing how this developmental block is bypassed by a gain-of-function mutation (T265I) in TaARF4-A2. This substitution targets an evolutionarily conserved residue within the DNA-binding domain, transforming the transcription factor into a hypermorphic repressor that unlocks latent developmental plasticity in the wheat culm. Crucially, this discovery defines the elusive molecular interface where spatial morphogens directly dictate temporal developmental clocks.

The T265I substitution offers a unique window into the evolutionary divergence of auxin signaling. While the loop region containing T265 is strictly conserved across land plants, our comparative analysis reveals a distinct functional dependency in wheat. Recent studies in basal land plants (*Marchantia polymorpha* and *Physcomitrium patens*) demonstrate that mutations in the flanking residues of this loop (corresponding to TaARF4-A2 T263 and M266) drive auxin insensitivity and enhance protein stability^34,35^. In contrast, our functional assays in hexaploid wheat demonstrate that substitutions at these flanking residues do not affect repressor activity. Instead, enhanced repression is specifically associated with alteration of the central T265 residue. Notably, a gain-of-function mutation at the corresponding position has been described in maize (*Truffula*, S281N), indicating that this central residue may represent a previously unrecognized hotspot for functional activation in grass ARFs. Our cross-species observations suggest that while the primary sequence of this regulatory ARF motif is deeply conserved, the specific amino acid dictating its functional state has diverged between basal lineages and advanced angiosperms.

Furthermore, this evolutionary divergence extends beyond the biochemical regulation of the ARF protein to the species-specific rewiring of its downstream transcriptional networks. At the physiological level, recent developmental studies in the basal land plant *M. polymorpha* reveal that the ancestral role of repressor B-ARFs (e.g., MpARF2) is to establish a localized, auxin-insensitive environment that protects meristematic pluripotency and governs primitive apical dominance^67,68^. As the NAP expanded and diversified during terrestrialization^28,30^, these orthologous junctions were likely repurposed. This explains the functional paradox where analogous central-residue mutations (e.g., S281N in maize versus T265I in wheat) yield remarkably divergent pleiotropic syndromes. In the maize lineage, ZmARF28 appears to be recruited primarily to regulate leaf initiation and floral sex determination. Conversely, in polyploid wheat, TaARF4-A2 evolved a specialized role to physically bridge the spatial and temporal networks, policing the SPL-mediated upper-aerial branching checkpoint. Consequently, the “locked” hyper-repressor in the *Bsh1* mutant triggers a systemic collapse of auxin-dependent phytohormonal homeostasis and directly represses *TaSPL14* and *TaSPL7* (Fig. 5 and Extended Data Fig. 10). This dual metabolic and transcriptional disruption creates a permissive environment for the ectopic outgrowth of upper nodes and bract derepression.

By establishing how an auxin-responsive factor directly governs the SPL timer, the TaARF4-2–SPL regulatory axis uncovered here provides a compelling example of how developmental heterochrony (an evolutionary shift in the timing of developmental events) can drastically reshape plant architecture. In wild-type wheat, the timely upregulation of *SPL* genes acts as a developmental clock, driving the transition to the reproductive phase and firmly restricting branching competence to the basal nodes (tillers). By acting as a hyper-repressor of *SPL* transcription, the Bsh-A1 variant delays this biological timer, effectively prolonging a “vegetative-like” state in the upper phytomers. This temporal decoupling permits upper AxMs to evade dormancy and initiate lateral outgrowth, a trait strictly suppressed during the domestication of major cereal crops. Thus, changes in developmental timing serve as a powerful mechanism to alter phytomer organ identity, highlighting how variations in the *SPL*-mediated molecular clock can drive the evolutionary diversification of shoot architectures across angiosperms.

From a breeding perspective, the TaARF4-2–SPL module provides a transformative framework for ideotype design. While the extreme aerial branching of *Bsh1* is agronomically unfavorable, it exposes a potent genetic lever. Rather than relying on severe structural mutations, fine-tuning this axis, perhaps by editing the upstream *cis*-regulatory elements of *TaARF4-2* to quantitatively modulate its expression, or by delicately tuning its downstream *SPL* targets, presents a strategy to optimize the trade-off between sink capacity and source potential. Similar to the successful engineering of the wheat *Q* gene^69,70^ and rice *RFL*^71^, manipulating such key junctions in conserved regulatory networks offers a promising avenue for unlocking yield potential in modern crops.

## Acknowledgements

We thank Prof. H. Mao and Z. Lu for generously sharing the *Taspl14&17-CR* sextuple knockout lines in the Fielder and ZM7698 backgrounds, respectively; Dr. Y Liu for providing chromatin accessibility data for the *TaSPL14* and *TaSPL7* homoeologs; Dr. D. Douchkov for technical assistance with serial image collection; L. Friedenberger for technical assistance with miRNA quantification; B. Kettig and D. Boehmert for technical assistance with the analysis of phytohormones; and Dr. A. Fiebig and D. Schüler for their help with data submission. This work was supported by the National Natural Science Foundation of China (grant number 32072054 to P.Q.), a China Scholarship Council (CSC) fellowship (no. 201906910089) to Z.G., and core funding from the Leibniz Institute of Plant Genetics and Crop Plant Research (IPK) to T.S. We also acknowledge the use of Gemini for English language editing and stylistic refinement during the manuscript preparation process.

## Contributions

Conceptualization: Z.G., P.Q., and T.S.; Methodology: Z.G. and Y.L.; Investigation: Z.G., Y.L., K.T., T.R., A.S., Q.C., Q.L., M.P., L.L., J.T., Y.A.T.M., M.K., S.Z., Y.H., R.F.H.G., N.W., J.K., Y.Z., Y.W., K.W., P.Q., and T.S.; Visualization: Z.G., Y.L., K.T., and S.O.; Funding acquisition: Z.G., P.Q., and T.S.; Supervision: Z.G., P.Q., and T.S.; Writing – original draft: Z.G.; review and editing: all authors.

## Data and materials availability

RNA-seq data are available from the European Nucleotide Archive (https://www.ebi.ac.uk/ena/browser/home) under the accession numbers PRJEB106556. All other data are available in the main text or the Extended Data materials. All materials are available upon request from the corresponding author.

## Materials and Methods

### Plant materials, growth conditions and phenotyping

#### Plant materials

The common wheat (*Triticum aestivum* L.) cultivar Shumai482 (SM482) was mutagenized with 0.6% ethyl methanesulfonate (EMS; Sigma-Aldrich). The original branching mutant line, designated *M4-15-1* (*Bsh1*), was isolated from a single heterozygous M_4_ individual. For genetic mapping, a segregating population was generated from a cross between *M4-15-1* and the wheat variety QZ212. A total of 132 individuals from a segregating F_4_ family were used for phenotypic segregation analysis. Bulked Segregant Exome-capture sequencing (BSE-seq) was performed using extreme phenotypic pools directly selected from this F_4_ generation. Recombinants were screened in the F_2:4_ to F_2:6_ generations for fine mapping. Based on genotype-phenotype co-segregation, two residual heterozygous lines (RHLs), RHL-1 and RHL-2, were identified. A pair of near-isogenic lines (NILs), designated *Bsh1* and WT, was subsequently developed from a single F_2:5_ heterozygous individual and used for physiological and molecular analyses.

Eleven additional branching mutants were identified from three independent EMS-mutagenized populations in the SM482, MM37, and Y26 genetic backgrounds. TILLING mutants in the Cadenza and Kronos backgrounds were obtained from SeedStor2 (https://www.seedstor.ac.uk/shopping-cart-tilling.php), and the Jing411 TILLING mutant was obtained from the Jing411 TILLING database^72^. The *TaARF4-2* triple loss-of-function mutant (*arf4-2_m1*) was generated by crossing three independent nonsense mutant lines followed by selfing. Taspl14/17 hextuple mutants (*Taspl14&17-CR*) were generated using CRISPR/Cas9 editing in the winter wheat variety ZM7698 and spring wheat variety Fielder as described in ^63,65,66^. Wheat variety Fielder was used for all stable transgenic transformations.

### Growth conditions

Field experiments: The F_2_ mapping population and selected segregating families from F_2:3_ to F_2:5_ generations were space-planted in a field at the Sichuan Agricultural University Experimental Station (Chengdu, Sichuan province, China) during the 2017 to 2022 growing seasons. Independent branching mutants were grown at the same location in the 2022-2023 growing season. Selected segregating families from F_2:6_ to F_2:7_ were space-planted in a field at the Leibniz Institute of Plant Genetics and Crop Plant Research (IPK) (Gatersleben, Germany) in 2023 and 2024, respectively.

Greenhouse and Chamber experiments: *N. benthamiana* plants were grown in a greenhouse at Sichuan Agriculture University under long-day conditions (16 h light/8 h dark) at 23 °C. T_0_ transgenic wheat plants were grown in the same greenhouse under 16 h of light at 24 °C and 8 h of dark at 16 °C. NILs, triple mutants, and transgenic plants (T_1_ and T_2_ generations) were grown in a greenhouse at IPK between 2022 and 2025 under long-day conditions (16 h light/8 h dark) at 19°/16°C (day/night). *Taspl14&17-CR* plants were grown in the same greenhouse in 2025. For transcriptomic and hormonal analyses requiring precise developmental staging, wheat plants were grown in a controlled phytochamber under conditions described by Thiel et al. (2021) (12 h light/12 h dark; 12°C/8°C, day/night). To synchronized germination for these experiments, seeds were germinated in 24-well trays and sampled directly without potting. For experiments requiring mature spike phenotyping, seeds were germinated in 96-well trays for two weeks, vernalized at 4°C for two weeks, acclimatized at 15°C for one week, and then transplanted into 11-cm pots until maturity.

*Arabidopsis* growth: *Arabidopsis thaliana* seeds were surface-sterilized and sown on half-strength Murashige and Skoog (½ MS) medium supplemented with 1% agar (pH 5.7). After stratification at at 4°C for 3 days in the dark, seedlings were grown at 22°C under a long-day conditions (16 h light/8 h dark) with a light intensity of 150 μmol m^−2^ s^−1^ photosynthetically active radiation.

### Phenotyping

All phenotypic data were collected from the main culm unless otherwise specified. Heading date was recorded when approximately half of the spike had emerged from the flag leaf sheath. Plant height was measured from the soil surface to the tip of the spike, excluding awns. Final spikelet number and rudimentary basal spikelet number were counted at the grain filling stage. In segregating populations, individual plants were classified as normal (wild type), branched (homozygous mutant) or intermediate (heterozygous), based on overall architecture and spike morphology.

### Culm node regeneration assay

Culm nodes subtending the spike (N1-N3) were harvested at the heading stage from wild-type and *Bsh1* plants, with the basal rachis node (N0) serving as a control. Explants were surface-sterilized by immersion in 70% (v/v) ethanol for 3 min, followed by incubation in 5% (v/v) sodium hypochlorite supplemented with 0.1% (v/v) Tween-20 for 15 min, and subsequently rinsed five times in sterile distilled water.

For *in vitro* culture, nodes were plated on K4NTS medium. This medium is a modified version of the K4NBT formulation described previously ^73^, in which 6-benzylaminopurine (BAP) and hygromycin were omitted, and maltose was substituted with 30 g/L sucrose. Plates were incubated at 21°C in the dark for 72 h, followed by transfer to a growth chamber at 24°C under a 16-h photoperiod (136 μmol s^−1^ m^−2^ photon flux density). After three weeks of regeneration, plantlets were transplanted into soil (13 cm pots) and acclimatized in a greenhouse.

### BSE-Seq

Two DNA bulks were constructed from the segregating F_4_ generation by pooling 27 individuals with the normal phenotype and 27 individuals with the extreme branched phenotype. Genomic DNA was extracted individually using a CTAB-based method, quantified, and pooled by mixing equal amounts of DNA from each individual.

Library preparation and sequencing were performed by Tcuni Technology (Chengdu, China). DNA quality was assessed using Qubit quantification and agarose gel electrophoresis. A total of 100 ng of pooled genomic DNA was utilized for library construction with the Twist Library Preparation EF Kit 2.0 (Twist Bioscience; 104207). The procedure included enzymatic fragmentation, end repair, A-tailing, and adapter ligation. Libraries were purified and size-selected using magnetic beads, amplified with indexed primers, and hybridized to an exome capture probe set at 65 °C for 16–24 h. Captured DNA fragments were recovered using streptavidin-coated magnetic beads, washed stringently, and eluted. Enriched libraries were subjected to limited-cycle PCR and quality controlled using Qubit and an Agilent Bioanalyzer before sequencing on a BGI DNBSEQ-T7 platform (paired-end 150 bp). Bulk segregant analysis (BSA) was performed according to previously established protocol^74^.

### Marker development and gene mapping

Single nucleotide polymorphisms (SNPs) identified from BSE-seq data were converted into Kompetitive Allele Specific PCR (KASP) or restriction enzyme-based CAPS/dCAPS markers. Using 132 individuals from the advanced segregating family (F_4_ generation) described above, the causal mutation was initially mapped to the long arm of chromosome 3A, within a genetic interval flanked by markers m3 and m11. Recombinants between these flanking markers were continuously screened from F_2:4_ to F_2:6_ generations. Combined genotypic and phenotypic analyses delimited the *Bsh1* locus to a narrower physical interval between markers m10 and m14, containing six high-confidence genes annotated in the wheat reference genome (IWGSC RefSeq v1.1). Marker m13 exhibited perfect co-segregation with the mutant phenotype across all identified recombinants. Primers used for map-based cloning are listed in Supplementary Table 1.

### Phylogenetic analysis

The TaARF4-A2 protein sequence (TraesCS3A02G449300) was used as a query against the Ensembl Plants database with default parameters to retrieve homologouos sequences from *Triticum aestivum*, *Hordeum vulgare*, *Brachypodium distachyon*, *Oryza sativa*, *Zea mays*, *Arabidopsis thaliana*, and *Physcomitrium patens*. Phylogenetic analysis was performed using amino acid sequences, and the maximum likelihood tree was generated using 1,000 bootstrap replicates.

### Scanning electron microscopy

Shoot apices containing immature spike and subtending culm nodes were collected from WT and *Bsh1* plants growing in the greenhouse at Waddington stages W2.5, W3, W3.5, W4, W5. Samples were fixed at 4°C in 50 mM cacodylate buffer (pH 7.2) containing 2% glutaraldehyde and 2% formaldehyde. After washing, samples were dehydrated in an ascending ethanol series, critical-point dried in a Bal-Tec critical point dryer (Leica microsystems, https://leica-microsystems.com), sputter-coated with gold in an Edwards S150B sputter coater (http://edwardsvacuum.com) and examined in a Zeiss Gemini30 scanning electron microscope (Carl Zeiss microscopy GmbH, https://zeiss.de) at 10 kV acceleration voltage. Recordings were saved as Tiff files.

### RNA extraction, qRT–PCR and RNA-seq

#### qRT–PCR

Total RNA was extracted using TRIzol reagent (Thermo Fisher Scientific) according to the manufacturer’s instructions. After genomic DNA removed prior to cDNA synthesis using the SuperScript III Reverse Transcriptase kit (Invitrogen; 18080044). Quantitative real-time PCR (qRT-PCR) was performed using Power SYBR Green PCR Master Mix (Applied Biosystems; 4367659) on a QuantStudio5 Real-Time PCR System (Thermo Fisher Scientific). Wheat *β-ACTIN* was used as the internal gene control. Relative transcript levels were calculated using the ΔΔCT method. Each experiment included three biological replicates. Primer sequences are listed in Supplementary Table 1.

#### miRNA qRT-PCR

Small RNAs were isolated using a miRNeasy Micro kit (Qiagen; 217084) and quantified with a Qubit Fluorimeter (Thermo Fisher Scientific). miRNA abundance was determined using the stem-loop qRT-PCR approach as previously described^75^.

Briefly, 500 ng of RNA was reverse transcribed using gene-specific stem-loop RT primers and the RevertAid First Strand cDNA Synthesis kit (Thermo Fisher Scientific; K1622). Quantitative PCR was performed using Power SYBR Green PCR Master Mix (Applied Biosystems; 4367659) on an Applied Biosystems 7900HT Fast Real-Time PCR system, following the manufacturer’s protocol. Data were analyzed in accordance with MIQE guidelines^76^. Relative expression levels were calculated using the ΔΔCT method and normalized to miR159 as an internal reference. Three biological replicates, each with three technical replicates, were analyzed.

### RNA-seq

Developing spikes and subtending upper node tissues (N1–N3) were carefully dissected from wild-type and *Bsh1* plants at the W2.5 (double ridge), W3 (glume primordium), and W3.5 (floret primordium) developmental stages. Total RNA was extracted using TRIzol reagent.

Poly-A libraries were prepared using TruSeq library preparation reagents (Illumina) and sequenced on a DNBSEQ-T7 platform (BGI). Clean reads were mapped to the wheat reference genome (IWGSC RefSeq v1.1) using TopHat2^77^. Differential gene expression analysis was performed using the DESeq2 R package. Genes with an adjusted *P* value (false discovery rate < 0.05) and |log_2_(fold change)| ≥ 1 were considered differentially expressed. Gene ontology enrichment analysis was conducted using the online tool Triticeae-GeneTribe^78^.

### Subcellular localization

Full-length *TaARF4-A2* coding sequences (CDS) was amplified from SM482 and *Bsh1* spikes, respectively, cloned into vector *pMD19-T* (TAKARA, number 3271), and subsequently subcloned into the transient expression vector *pJIT163* to generate BSH-A1–eGFP (WT) or Bsh-A1–eGFP (mutant) fusions. Wheat protoplasts were isolated from the first leaf of 7-day-old seedlings and transformed as described previously (Yoo et al., 2007). Fluorescence signals were examined 16 h after transformation using a confocal microscope (LSM780, Zeiss). Primers are listed in Supplementary Table 1.

### Plant transformation

*TaARF4-A2* CDS from SM482 (T265) and *Bsh1* (I265) were cloned into the *pWMB003* vector (*ZmUbi* promoter, Nos Terminator) to generate intermediate constructs. The expression cassette was transferred into *pWMB111* (containing the Bar selection cassette driven by *ZmUbi*) via HindIII digestion and ligation to generate the target expression vectors *pWMB111-TaARF4-A2* (T265) and *pWMB111-TaARF4-A2* (I265). Final constructs were used for *Agrobacterium*-mediated transformation of wheat variety fielder.

For *Arabidopsis* transformation, *TaARF4-A2* CDS from SM482 and *Bsh1* were cloned into *TSK108* fused with tandem FLAG tags and transferred to *pB7WG2* (35S promoter) by Gateway cloning (Invitrogen; 12535029). Constructs were introduced into *Arabidopsis* Col-0 by floral dipping method. Two independent homozygous lines for each construct were selected with phosphinothricin. Flowering time was assessed by counting rosette and cauline leaves when the first floral buds became visible. Primers for are listed in Supplementary Table 1.

### mRNA *in situ* hybridization

Gene-specific fragments (∼265 bp) were amplified from SM482 spike cDNA and cloned into *pGEM-T* cloning vector. Sense (negative control) and antisense probes were generated using *in vitro* transcription with T7 RNA polymerase. Templates were produced using fusion primers containing a 20 bp T7 promoter sequence (5′-TAATACGACTCACTATAGGG-3′).

Shoot apices were dissected and fixed overnight in FAA (50% ethanol, 5% acetic acid, and 3.7% formaldehyde) at 4°C, dehydrated through an ethanol series (50, 70, 85, 95, and 100%), and embedded in Paraplast Plus (Kendall). Sections (8 μm thick) were prepared, mounted onto Superfrost plus slides, and hybridization was performed as described previously^79^. Primers are listed in Supplementary Table 1.

### Light microscopy and 3D reconstruction

Samples were fixed with 2% glutaraldehyde and 2% formaldehyde in 50 mM cacodylate buffer (pH 7.2). After dehydration probes were embedded in Spurr resin followed by serial sectioning on a Reichert-Jung Ultracut S (Leica, Vienna, Austria) using section thickness of 1 µm and 10 µm interval. After staining with aqueous kristal-violet dye sections were recorded in a Zeiss Axio Scan Z1 slidereader (Carl Zeiss, Oberkochen, Germany) using a 10x NA 0.45 objective. Following contrast and brightness adjustment in Adobe Photoshop (version 25.12.4), image stacks were aligned with ImageJ open source software (version 1.54m). Segmentation of vascular structures and 3D reconstruction was performed with Amira software (Thermo Fischer Scientific, version 6.5.0).

### Phytohormone quantification

For phytohormone measurements (IAA, iPR, and tZR), culm nodes at approximately Waddington stage W8.0 were collected from WT and *Bsh1* plants. Each sample contained four to five biological replicates, with each replicate consisting of pooled tissue from five nodes. Samples were immediately frozen in liquid nitrogen and stored at −80°C. Hormones were quantified using UPLC-MS platform at IPK (Gatersleben, Germany) following published procedures^80^.

### Agar-based and hydroponic assays

Agar plates (2%) containing IAA at 0, 10^−8^ M, 10^−7^ M, 10^−6^ M, 10^−5^ M, or 10^−4^ M were prepared in 13-cm square Petri dishes. The wheat seeds were germinated on wet filter papers and transferred to agar plates. Seedlings were laid horizontally for half day and then plates were placed upright in a climate chamber under long-day conditions (16 h light/8 h dark) at 20°C /18°C (day/night), 70% humidity and a light intensity of 250 μmol m^−2^ s^−1^. The longest seminar root was measured daily.

Hydroponics were performed in the same climate chamber. Seedlings were transferred to full nutrient solution 4 days after germination. The nutrient solution contained 2 mM Ca(NO_3_)_2_, 0.5 mM K_2_SO_4_, 0.5 mM MgSO_4_, 0.1 mM KH_2_PO_4_, 0.1 mM KCl, 1 μM H_3_BO_3_, 0.5 μM MnSO_4_, 0.5 μM ZnSO_4_, 0.2 μM CuSO_4_, 0.01 μM (NH_4_)_6_Mo_7_O_24_ and 0.1 mM Fe-EDTA. After 3 days, the nutrient solution was renewed and IAA (100 mM stock in DMSO) was added to a final concentration of 10 μM. DMSO was used as a mock control. After 4 days of treatment, roots were harvested, fixed in 75% ethanol, scanned, and analysed using WinRHIZO Pro (Regent Instruments).

For qRT-PCR, roots were harvested at 0 h, 0.5 h, and 3 h after treatment, immediately frozen in liquid nitrogen and stored at -80°C.

### IAA and NPA soil treatments

IAA and NPA treatment assays were performed described previously^81^. Stock solutions for indole-3-acetic acid (IAA; Sigma-Aldrich; I3750) and N-1-naphthylphthalamic acid (NPA; MedChemExpress; HY-116425) were separately prepared in 95% ethanol and diluted in water containing 0.02% Silwet L-77 (PlantMedia) before application. Treatments were performed by applying 50 ml of working solution to WT and *Bsh1* plants grown in 9 cm^2^ pots. Ethanol diluted solution (mock) was used as a control. Treatments were applied daily from the double ridge stage for 4 weeks and subsequently every 2 days until the end of the experiment.

### EMSA

The full-length CDS of *TaARF4-A2* from SM482 (T265) and *Bsh1* (I265) were cloned into the *pMAL-C2x* vector with a fusion with maltose-binding protein (MBP), separately. Then they were transformed into E. coli strain BL21 (DE3) and purified *in vitro* by Amylose Resin (New England Biolabs; E8021). For EMSA, we synthesized short fragment, 26 bp and 32 bp, containing the TGTCGC motif from *TaSPL-D14* and *TaSPL-D7* promoters, respectively, which were labelled with Cyanine 5.5 (Cy5.5) at the 5’ end (Eurofins). We used an unlabeled probe for competition assays. The binding reaction was performed in 20 μL volumes using the Odyssey® glycerol at room temperature for 30 min. 6% native polyacrylamide gels in 0.5× TBE was used to sperate the DNA-protein complexes, which were later analyzed using the LI-COR Odyssey Imaging System. All primers and probe sequences used for EMSA are shown in Supplementary Table 1.

### Dual-luciferase transcriptional activity assays

Effector constructs were generated by cloning the full-length CDS of *TaARF4-A2* from SM482 (T265) and *Bsh1* (I265) into the *pCAMBIA1300*-mScarlet3 or *pCAMBIA1300*-eGFP vectors driven by the CaMV 35S promoter. An empty vector served as the negative control. To assess the impact of key residue substitutions, site-directed mutagenesis (T265N, T263K, and M266Q) was performed using specific primers. The 2 kb full-length promoter region of *TaSPL-D14* was amplified from the SM482 genome and cloned into the pGreenII-0800-LUC vector, which contains the firefly luciferase (*LUC*) gene as the reporter and the *Renilla* luciferase (*REN*) gene as an internal control.

All effector and reporter plasmids were transformed into *Agrobacterium tumefaciens* strain GV3101 carrying the p19 helper plasmid to suppress post-transcriptional gene silencing. *Nicotiana benthamiana* leaves were co-infiltrated with the bacterial suspensions. Two days post-infiltration, leaf discs were harvested and homogenized in liquid nitrogen. Firefly and *Renilla* luciferase activities were quantified using the Dual-Luciferase Reporter Assay System (Promega). Relative luciferase activity (LUC/REN) was calculated to normalize transformation efficiency. Experiments were performed with three to six biological replicates.

### Statistical analysis

Statistical analyses and data visualization were performed using GraphPad Prism v.10 (GraphPad Software) and R software. In bar graphs, data are presented as mean ± s.d., with individual data points overlaid for each biological replicate. For comparisons between two groups, statistical significance was determined using a two-sided Student’s t-test. For multiple group comparisons, a one-way analysis of variance (ANOVA) was performed, followed by Tukey’s honestly significant difference post-hoc analysis. Different letters indicate statistically significant differences (*P* < 0.05). Specific sample sizes (n) and detailed statistical parameters are provided in the respective figure legends.

## Supplementary Information

**Supplementary Fig. 1:** Phenotypic characterization of the original *Bsh1* mutant line *M4-15-1* and the developmental workflow of genetic populations.

**Supplementary Fig. 2:** Extended phenotypic and agronomic characterization of *Bsh1*.

**Supplementary Fig. 3:** Histological characterization of ectopic meristems and regenerative competency of upper culm nodes in *Bsh1*.

**Supplementary Fig. 4:** Spatial expression landscapes of *ARF4* and *SPL* orthologs in tetraploid wheat spikes.

**Supplementary Fig. 5:** Structural predictions of wild-type and mutant TaARF4 DNA-binding domains.

**Supplementary Fig. 6:** Auxin sensitivity assays and pharmacological treatment experiments.

**Supplementary Fig. 7:** RNA-seq experimental design and global transcriptome profiling.

**Supplementary Fig. 8:** Expression profiles of key DEGs involved in hormone signaling and developmental regulation.

**Supplementary Table 1:** Primer sequences used in this study.

**Supplementary Video 1: 3D spatial reconstruction and rotational overview of the vascular network.** Three-dimensional rotational visualization corresponding to the node architecture in Fig. 4f, illustrating the overall spatial topology and organization of the vascular bundles.

**Supplementary Video 2: Sequential z-axis assembly of the color-coded 3D vascular topology.** Bottom-to-top dynamic assembly of transverse sections. Individual vascular bundles are color-coded to track their continuity and spatial distribution across the vertical axis of the node.

**Extended Data Fig. 1:**
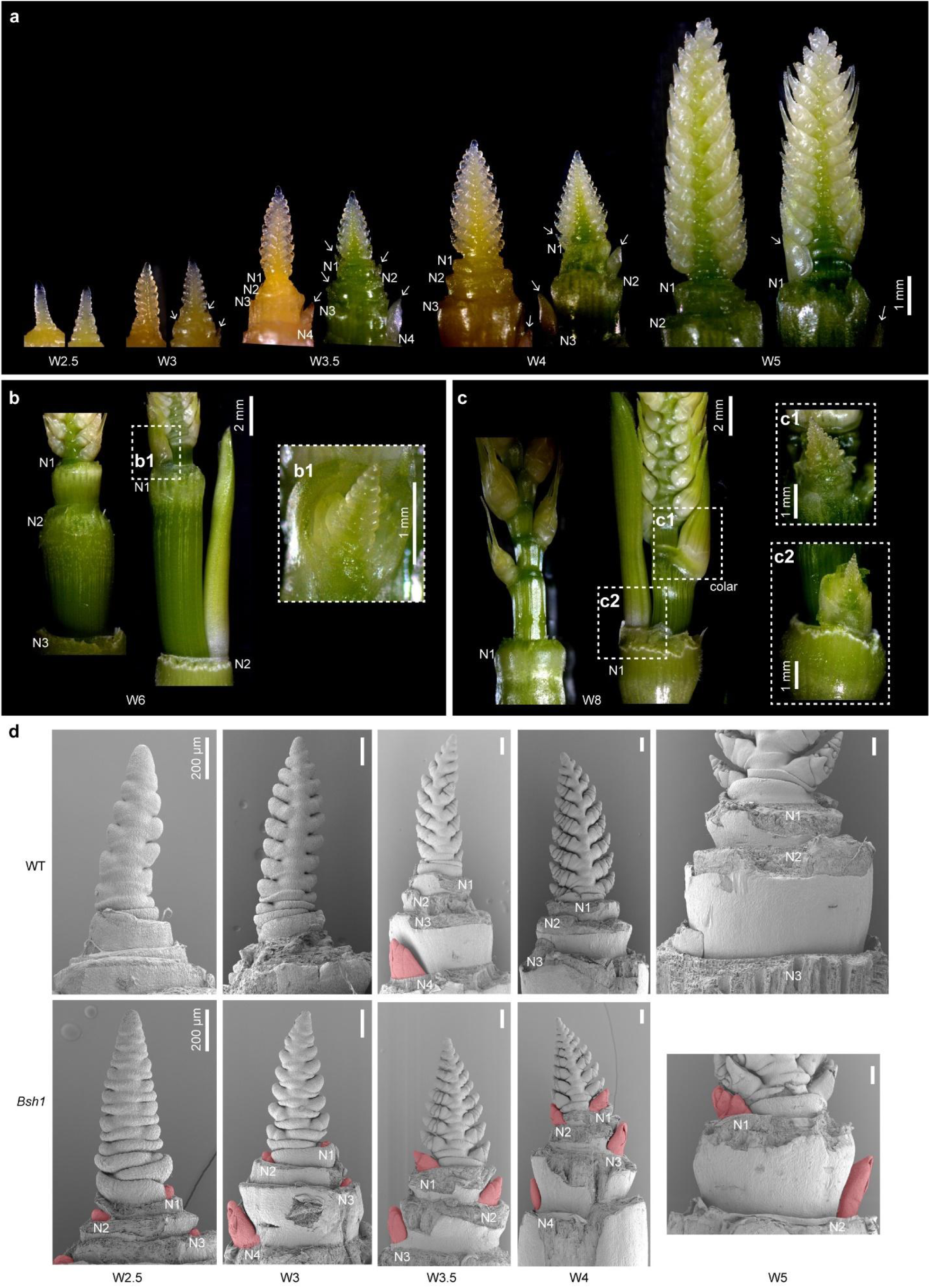
Developmental time-course of ectopic axillary meristem initiation in *Bsh1.* **a–c**, Stereomicroscopic tracking of axillary bud development in *Bsh1* across different stages of spike differentiation. (WT on the left; *Bsh1* on the right). **a**, Representative shoot apices from the double ridge stage (W2.5) to the awn primordium stage (W5). White arrows indicate axillary buds initiated at the upper nodes (N1–N4) of the main culm, which remain dormant in WT (typically N1-N3). **b**,**c**, Continued development of these ectopic buds at W6 (**b**) and W8 (**c**). Dashed boxes indicate regions enlarged in (**b1**), (**c1**), and (**c2**), showing that ectopic buds differentiate into multiple leaves and young spikes. **d**, Comparative scanning electron microscopy (SEM) analysis of axillary meristem (AxM) initiation. WT shoot apices (top) show smooth node surfaces (N1–N3) below the developing spike, with no detectable AxM initiation from W2.5 to W5. In contrast, *Bsh1* shoot apices (bottom) display ectopic AxM initiation at the corresponding nodes (pseudo-colored in red). Ectopic AxMs are detectable as early as W2.5 and progressively enlarge. Scale bars are indicated in the panels. Developmental stages are defined according to the Waddington scale (W). N1 denotes the first node below the spike.

**Extended Data Fig. 2:**
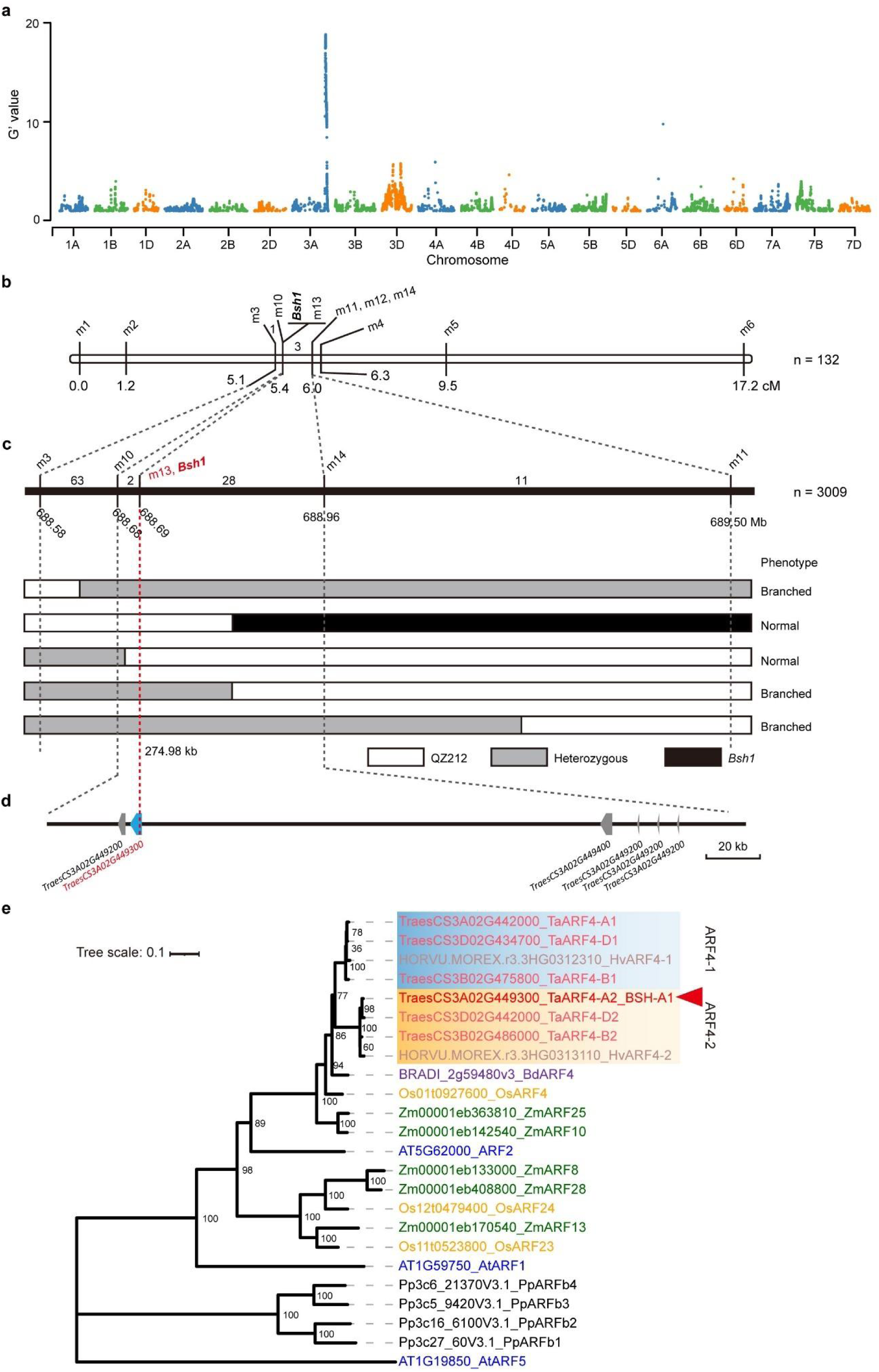
Map-based cloning and phylogenetic analysis of *TaARF4-A2*. **a,** Genome-wide linkage analysis using the G’ statistic derived from bulked-segregant exome sequencing (BSE-Seq) of extreme phenotypic pools selected from the segregating F_4_ generation. A major peak on the long arm of chromosome 3A identifies the candidate region harboring the *Bsh1* locus. **b,** Initial genetic linkage map constructed using 132 individuals from the advanced segregating family (F_4_ generation), delimiting the *Bsh1* locus between markers m3 and m4. Genetic distances are given in centiMorgans (cM). **c,** High-resolution fine-mapping of *Bsh1*. Screening of 3,009 individuals from the advanced F_2:5_ and F_2:6_ generations narrowed the locus to a 274.98-kb physical interval flanked by markers m10 and m14. The mutant phenotype completely co-segregates with marker m13. Graphical genotypes of key recombinants are shown (white, wild-type parent QZ212; grey, heterozygous; black, *Bsh1* mutant parent). Corresponding phenotypes (Branched or Normal) are indicated. Physical coordinates are based on the Chinese Spring Reference Genome (IWGSC RefSeq v1.1). **d,** Gene annotation within the 275-kb candidate interval. Six high-confidence protein-coding genes predicted (grey arrows). The causal gene, *TraesCS3A02G449300* (*TaARF4-A2*), is highlighted in blue. **e,** Maximum-likelihood phylogenetic tree of ARF4 proteins from representative plant species. Bootstrap values (from 1,000 replicates) are indicated at the nodes. TaARF4-A2 (BSH-A1; red arrowhead) clusters with the monocot ARF4 clade and is the direct ortholog of Arabidopsis *AtARF2* and rice *OsARF4*. Species abbreviations: Ta, *Triticum aestivum*; Hv, *Hordeum vulgare*; Bd, *Brachypodium distachyon*; Os, *Oryza sativa*; Zm, *Zea mays*; At, *Arabidopsis thaliana*; Pp, *Physcomitrium patens*.

**Extended Data Fig. 3:**
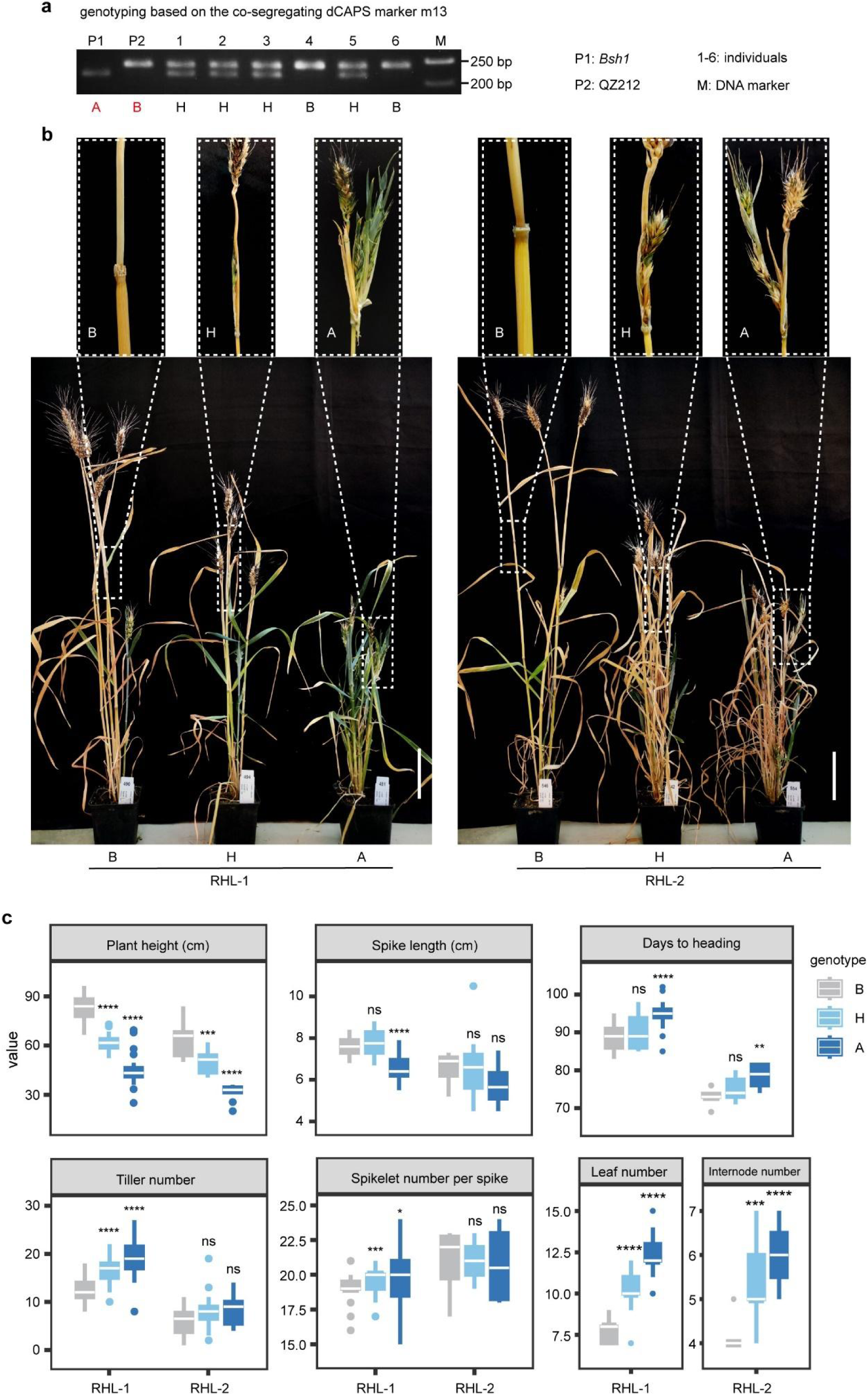
Validation of the *Bsh1* locus using residually heterozygous lines (RHLs). **a,** Genotyping using the derived cleaved amplified polymorphic sequence (dCAPS) marker m13, which co-segregates with the *Bsh1* phenotype. P1, *Bsh1* parent (mutant allele); P2, QZ212 parent (WT allele). A, homozygous mutant; B, homozygous WT; H, heterozygous. M, DNA marker. **b,** Co-segregation of phenotype with marker m13 in two independent RHL populations (RHL-1 and RHL-2). Plants with genotype A recapitulate the *Bsh1* syndrome (dwarfism, aerial branching, spike deformities), B plants display WT architecture, and H plants show an intermediate semi-dominant phenotype. Insets show magnified views of node and spike morphology. Scale bars, 10 cm. **c,** Agronomic traits comparison among genotypes (A, B, H) within RHL-1 and RHL-2. Traits include plant height, spike length, days to heading, tiller number, spikelet number per spike, leaf number, and internode number. Box plots show the median (center line), interquartile range (box), and 1.5× the interquartile range (whiskers). P values were calculated by two-sided Student’s *t*-test relative to genotype B. \**P* < 0.05; \*\**P* < 0.01; \*\*\**P* < 0.001; \*\*\*\**P* < 0.0001; ns, not significant.

**Extended Data Fig. 4:**
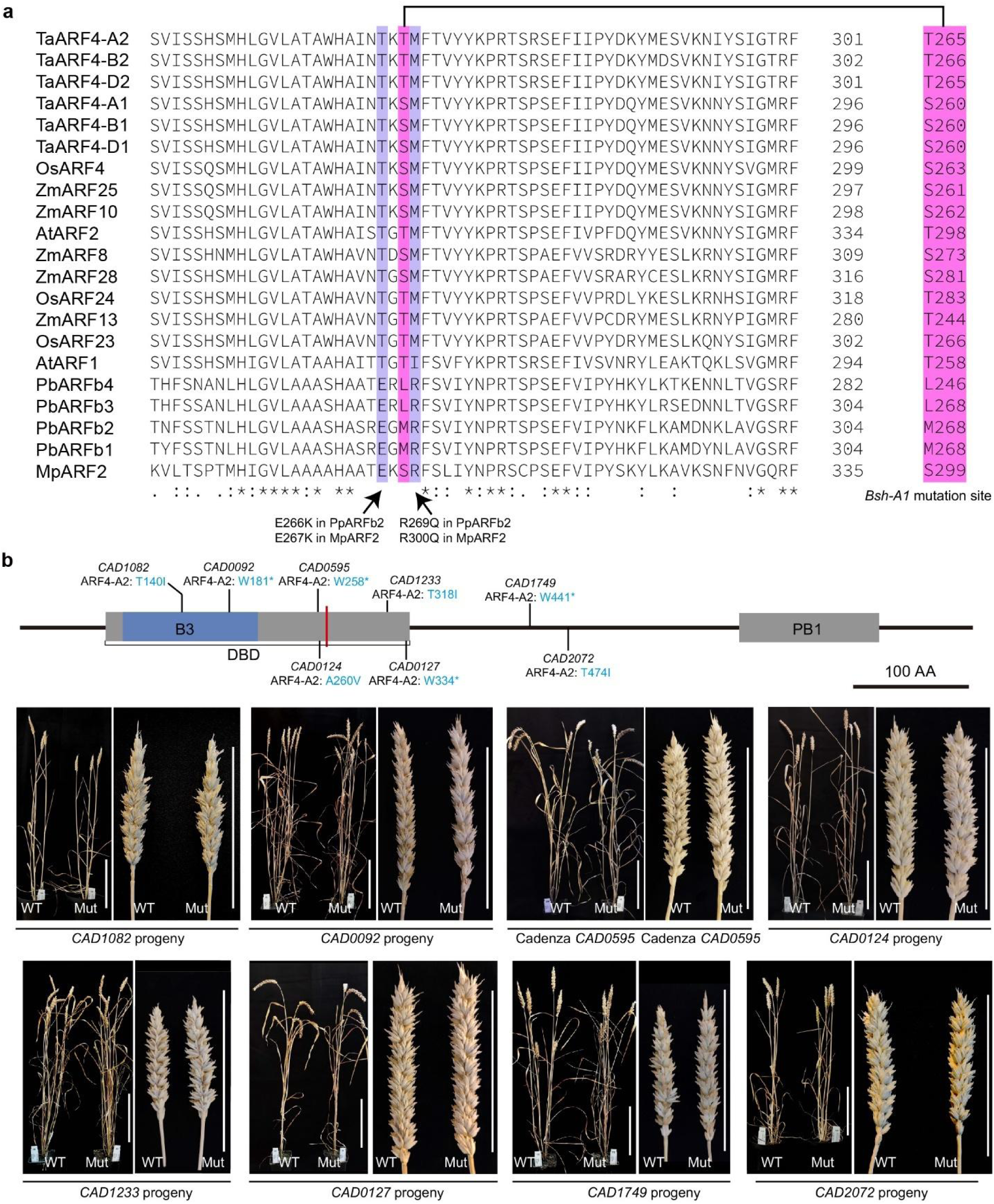
Evolutionary conservation of the T265 residue and phenotypic characterization of *TaARF4-A2* TILLING mutants. **a,** Multiple sequence alignment of the loop region within the DNA-binding domain of ARF4 orthologs from wheat (*Ta*), rice (*Os*), maize (*Zm*), *Arabidopsis* (*At*), *Physcomitrium patens* (*Pp*), and *Marchantia polymorpha* (*Mp*). The conserved T265 residue mutated in *Bsh1* is highlighted (pink). Adjacent residues (purple) correspond to sites reported to affect protein stability in bryophytes (e.g., E267K and R300Q in *MpARF2*). **b,** Identification and phenotypic analysis of eight independent *TaARF4-A2* TILLING mutants in hexaploid wheat cv. Cadenza. Top, gene model and protein domain (B3 DNA-binding domain and PB1 domain) with mutation positions. Four nonsense alleles (*CAD0092*, *CAD0595*, *CAD1233*, *CAD1749*) and four missense alleles (*CAD1082*, *CAD0124*, *CAD0127*, *CAD2072*) are shown; asterisks indicate stop codons. Bottom, representative whole-plant and spike phenotypes of homozygous mutants (Mut) and WT segregants (WT). None of these loss-of-function alleles recapitulate the *Bsh1* aerial-branching or spike-deformity phenotypes. Scale bars, 10 cm.

**Extended Data Fig. 5:**
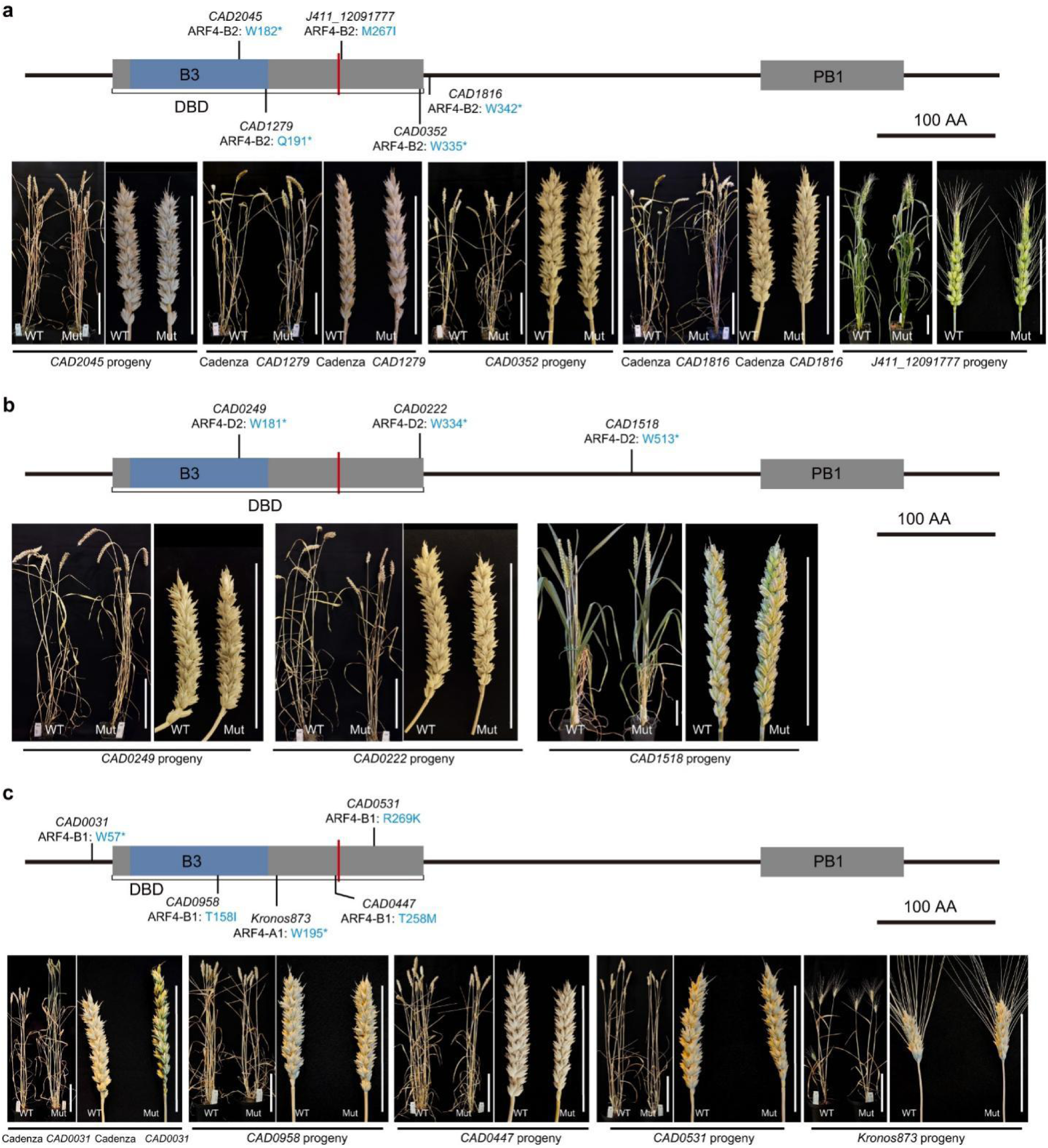
Phenotypic characterization of TILLING mutants in *TaARF4* homoeologs and paralogs. **a–c,** Identification and phenotypic analysis of TILLING mutants targeting *TaARF4-A2* homoeologs and paralogs. **a,** *TaARF4-B2* mutants. Top, gene model showing four nonsense alleles (*CAD2045*, *CAD1279*, *CAD0352*, *CAD1816*) and one missense allele (*J411_12091777*, M267I). M267I corresponds to the residue adjacent to the T265-equivalent site and overlaps with a stability-related residue described in Marchantia. Bottom, representative phenotypes of homozygous mutants and WT segregants. **b,** *TaARF4-D2* mutants. Top, gene model showing three nonsense alleles (*CAD0249*, *CAD0222*, *CAD1518*). Bottom, representative phenotypes of homozygous mutants and WT. **c,** Paralogs *TaARF4-B1* and *TaARF4-A1*. Top, gene model showing one nonsense allele (*CAD0031*) and three missense alleles (*CAD0958*, *CAD0447*, *CAD0531*) in *TaARF4-B1*, and one nonsense allele in *TaARF4-A1* (*Kronos873*). Mutations include a missense allele (*CAD0447*; T258M, corresponding to the stability-related "E" site). Bottom: representative phenotypes. For all panels, gene models indicate the B3 DNA-binding domain (blue) and PB1 domain (grey). Asterisks indicate stop codons. Neither nonsense alleles nor specific missense alleles at conserved adjacent residues (M267I, T258M) phenocopy *Bsh1*. Scale bars, 10 cm.

**Extended Data Fig. 6:**
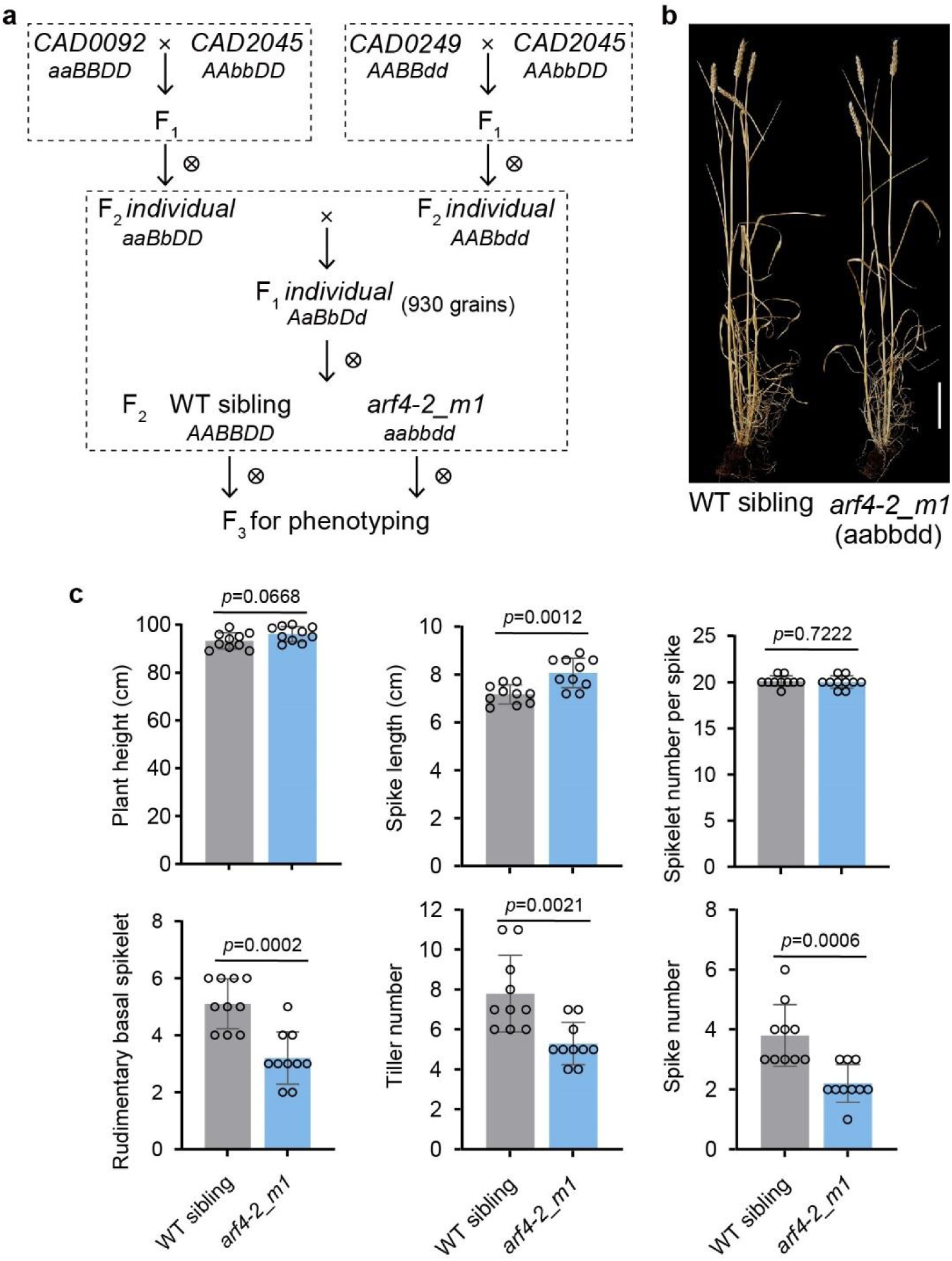
Generation and characterization of a *TaARF4-2* triple loss-of-function mutant. **a,** Crossing scheme used to generate the triple loss-of-function mutant *arf4-2_m1* (*aabbdd*). Independent nonsense mutants in *TaARF4-2* A (*CAD0092*), B (*CAD2045*), and D (*CAD0249*) homoeologs were crossed and selfed. F_2_ plants were genotyped to identify the triple homozygous mutant (*aabbdd*) and the corresponding WT sibling (*AABBDD*). F_3_ progeny were used for phenotyping. **b,** Representative mature phenotypes of *arf4-2_m1* and its WT sibling. The triple mutant shows reduced tillering and increased height but lacks ectopic aerial branching. Scale bar, 10 cm. **c,** Agronomic traits comparison between WT sibling and *arf4-2_m1*. Traits evaluated include plant height, spike length, spikelet number per spike, rudimentary basal spikelet number, tiller number, and spike number. Data are means ± s.d. (n = 10). *P* values were determined by two-sided Student’s *t*-test.

**Extended Data Fig. 7:**
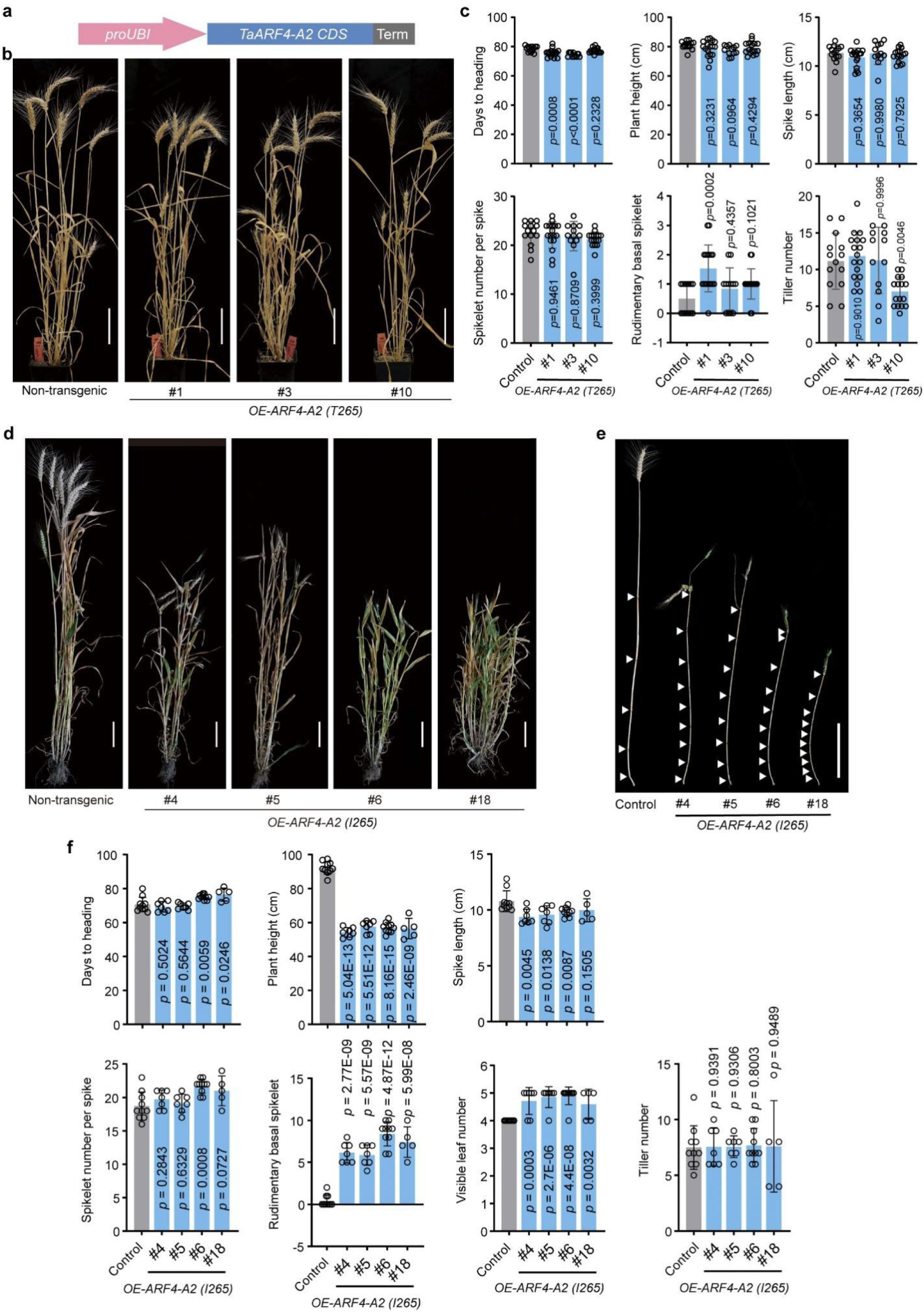
Phenotypic analysis of transgenic wheat lines overexpressing WT or *Bsh1 TaARF4-A2* alleles. **a,** Schematic of the transformation construct. The *TaARF4-A2* coding sequence (WT T265 or mutant I265) was driven by the maize Ubiquitin promoter (*proUBI*). The WT CDS was fused with an HA-tag. **b,c,** Characterization of independent lines overexpressing the WT allele (*OE-ARF4-A2* T265). **b,** Representative whole-plant phenotypes of non-transgenic controls and three independent OE lines (#1, #3, #10) at maturity. Overexpression of WT *TaARF4-A2* does not induce aerial branching or major architectural defects. Scale bars, 10 cm. **c,** Agronomic traits quantification for WT overexpression lines. Data are means ± s.d. (n ≥10). *P* values were determined by two-sided Student’s *t*-test comparing each transgenic line to the control. **d–f,** Characterization of independent lines overexpressing the mutant allele (*OE-ARF4-A2* I265). **d,** Representative whole-plant phenotypes of controls and four independent lines (#4, #5, #6, #18), which recapitulate the *Bsh1* syndrome. Scale bars, 10 cm. **e,** Main culms with leaf sheaths removed, revealing increased node number (white arrowheads). Scale bar, 10 cm. **f,** Agronomic traits quantification for I265 overexpression lines. Data are means ± s.d. (n ≥ 5). *P* values were determined by two-sided Student’s *t*-test.

**Extended Data Fig. 8:**
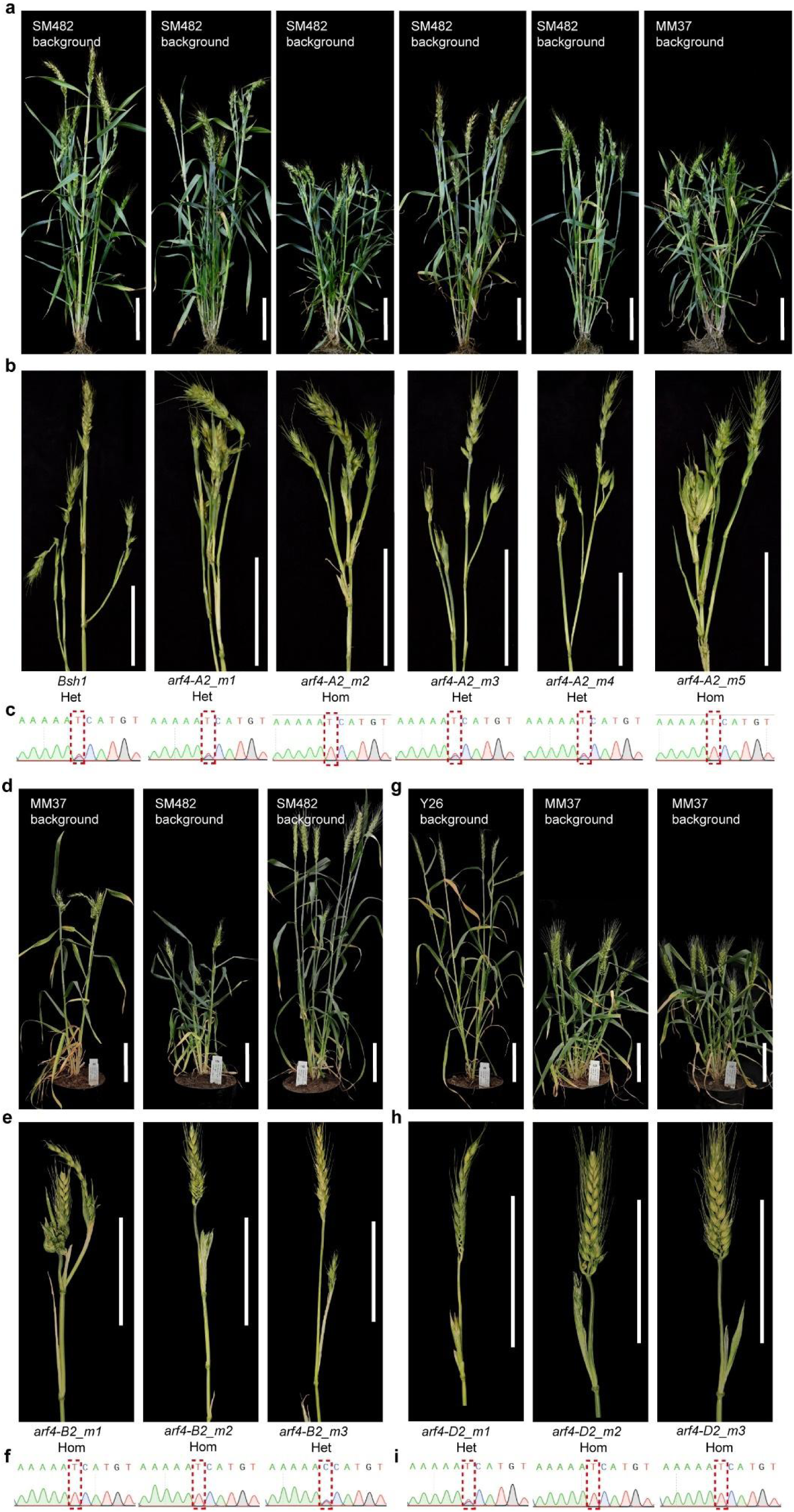
The T265I substitution confers a dominant hypermorphic phenotype across all three *TaARF4-2* homoeologs. **a-c,** Characterization of five independent *TaARF4-A2* mutant alleles (*arf4-A2_m1* to *arf4-A2_m5*) identified from EMS-mutagenized populations (SM482 or MM37 backgrounds) that carry the identical T265I substitution as the original *Bsh1* mutant. **a,** Representative whole-plant phenotypes exhibiting dwarfism and increased tillering. **b,** Close-up views of spikes and upper culm nodes. Zygosity is indicated (Het, heterozygous; Hom, homozygous). All alleles perfectly recapitulate the ectopic aerial branching and spike deformities characteristic of *Bsh1*. **c,** Sanger sequencing chromatograms confirming the recurrent C-to-T transition. **d–f,** Characterization of three independent *TaARF4-B2* mutants (*arf4-B2_m1* to *m3*). **d,** Whole-plant phenotypes (MM37 or SM482 backgrounds). **e,** Close-up views of spikes and nodes. **f,** Sanger chromatograms showing the C-to-T transition leading to the orthologous T-to-I substitution. **g–i,** Characterization of three independent *TaARF4-D2* mutants (*arf4-D2_m1* to *m3*). **g,** Whole-plant phenotypes (Y26 or MM37 backgrounds). **h,** Close-up views of spikes and nodes. **i,** Sanger chromatograms showing the corresponding C-to-T transition in *TaARF4-D2*. Collectively, these independent recurrent mutations demonstrate that the T265I substitution robustly rewires plant architecture and releases axillary bud dormancy regardless of whether it in the A, B, or D subgenome. Scale bars, 10 cm (**a, b, d, e, g, h**).

**Extended Data Fig. 9:**
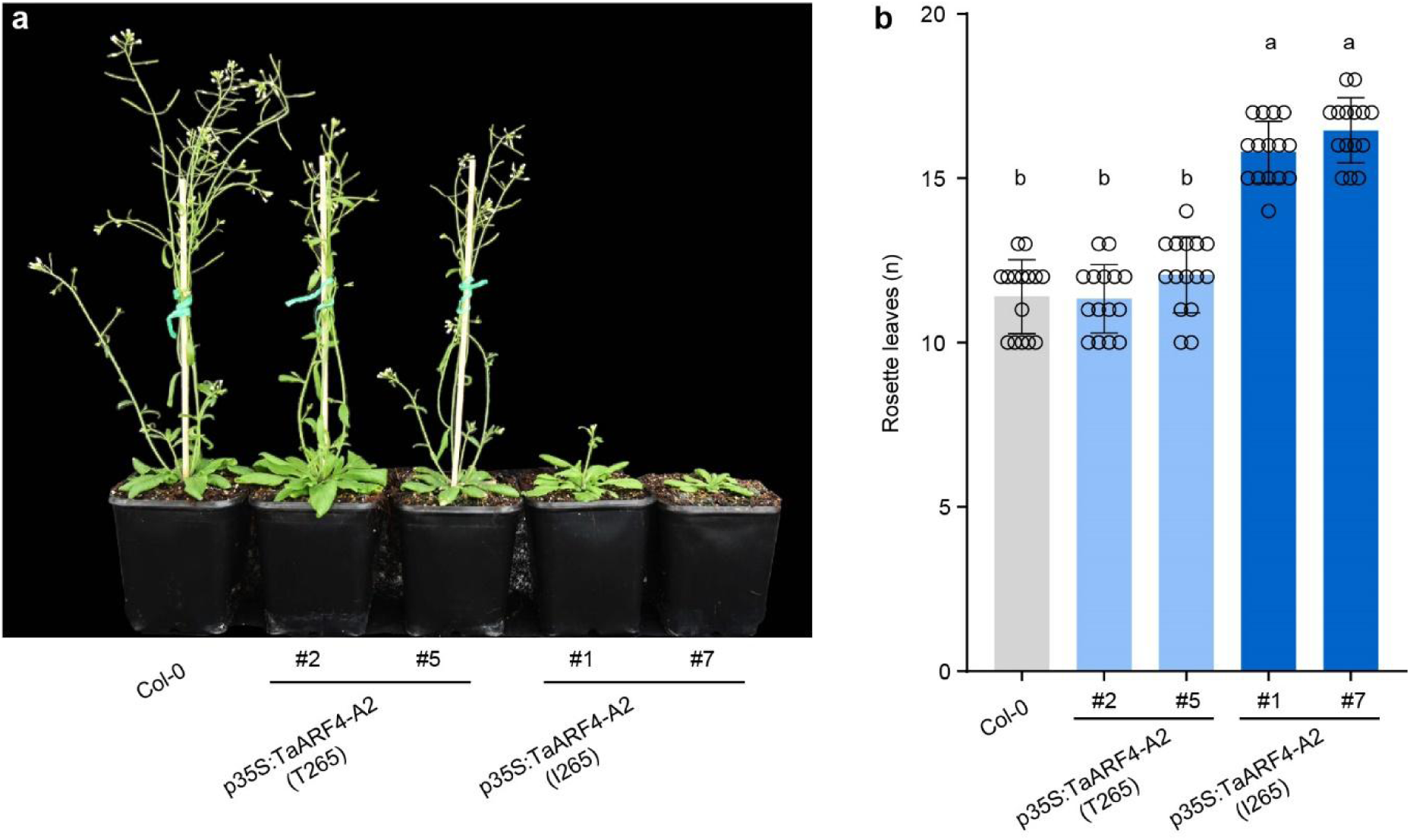
Functional conservation of the Bsh-A1 hypermorphic activity in *Arabidopsis*. **a,** Representative phenotypes of *Arabidopsis* Col-0 and heterologous overexpression lines expressing either the wild-type wheat *TaARF4-A2* (*p35S:TaARF4-A2*_T265_; lines #2 and #5) or the mutant allele (*p35S:TaARF4-A2*_I265_; lines #1 and #7) at the bolting stage (∼ 5 weeks). The I265 lines exhibit delayed bolting and reduced inflorescence development compared to Col-0 and T265 overexpression lines. **b,** Flowering time quantification as rosette leaf number at bolting (n = 15 biologically independent plants per line). The I265 overexpression lines produced significantly more rosette leaves before transitioning to the reproductive phase, indicating a prolonged vegetative stage. Data are means ± s.d. Different letters indicate significant differences (*P* < 0.05) as determined by one-way ANOVA with Tukey’s multiple comparisons test. These results demonstrate that the T265I substitution confers a conserved hyper-repressive activity to TaARF4-A2, effectively delaying the vegetative-to-reproductive phase transition in a heterologous dicot system.

**Extended Data Fig. 10:**
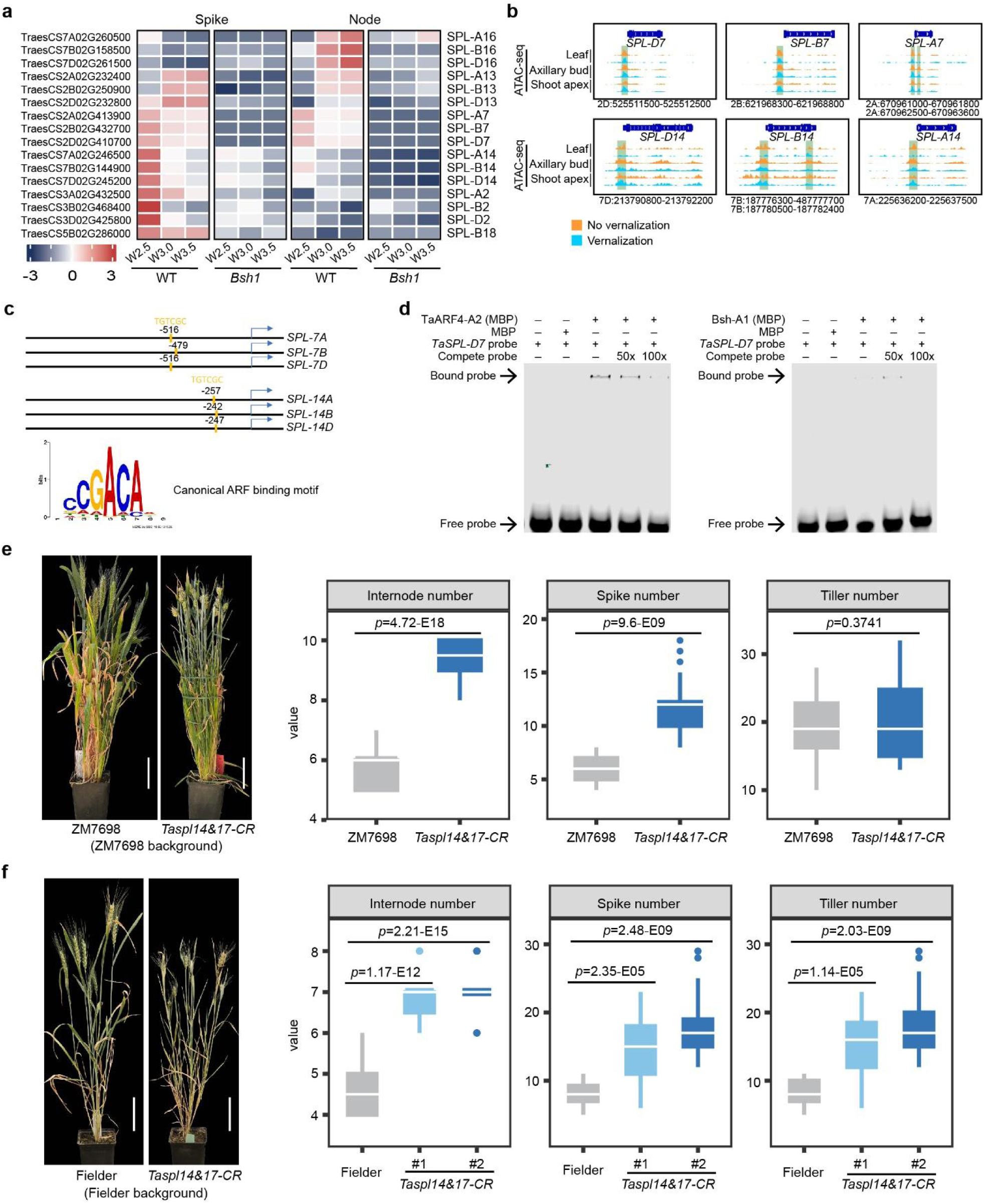
Repression of *TaSPL* targets and phenotypic characterization of *Taspl14/17* sextuple mutants. **a,** Heatmap of *SPL* family gene expression (row-scaled Z-scores) in WT and *Bsh1* developing spikes and nodes across successive developmental stages. The data highlight the broad transcriptional repression of multiple *SPL* homoeologs in *Bsh1*, including *TaSPL7*, *TaSPL13*, and *TaSPL14*. **b,** Chromatin accessibility at *TaSPL7* and *TaSPL14* promoter regions based on published ATAC-seq datasets. Tracks display open chromatin peaks in leaf, axillary bud, and shoot apex tissues under vernalized (blue) and non-vernalized (orange) conditions, indicating that these regulatory regions are highly accessible for transcription factor binding. Green boxes indicate accessible regions containing AuxRE motifs identified in (**c**). **c,** Schematic representation of *TaSPL7* and *TaSPL14* homoeologous promoters (A, B, and D subgenomes). Canonical ARF-binding motifs (AuxRE: TGTCGC) are mapped relative to the transcription start site (TSS). A sequence logo of the highly conserved AuxRE motif is shown below. **d,** Electrophoretic mobility shift assay (EMSA) demonstrating that both wild-type TaARF4-A2 (MBP) and *Bsh1* mutant (MBP-Bsh1-A1) proteins retain the intrinsic capacity to directly recognize and bind the canonical auxin response element within the target *TaSPL-D7* promoter *in vitro*. Unlabeled competitor probes (50× and 100×) were added to confirm binding specificity. (+) and (−) indicate the presence or absence of respective components. Black arrows indicate the bound protein-DNA complexes (shifted bands) and free probes. **e, f,** Whole-plant phenotypes (left) and agronomic quantification (right; internode, spike, and tiller numbers) of the *Taspl14&17* CRISPR sextuple mutants (*Taspl14&17-CR*) in the winter wheat cv. ZM7698 (**e**) and spring wheat cv. Fielder (**f,** showing two independent lines) backgrounds. Detailed spike and nodal branching phenotypes are presented in Fig. 5h. For the box plots, the center line represents the median, box limits indicate upper and lower quartiles, and whiskers indicate the data range. *P* values were determined by two-sided Student’s *t*-test. (n ≥10 biologically independent plants).

