## Supplementary figures for "Hypermorphic TaARF4 shapes wheat architecture by repressing SPL-mediated developmental timing"

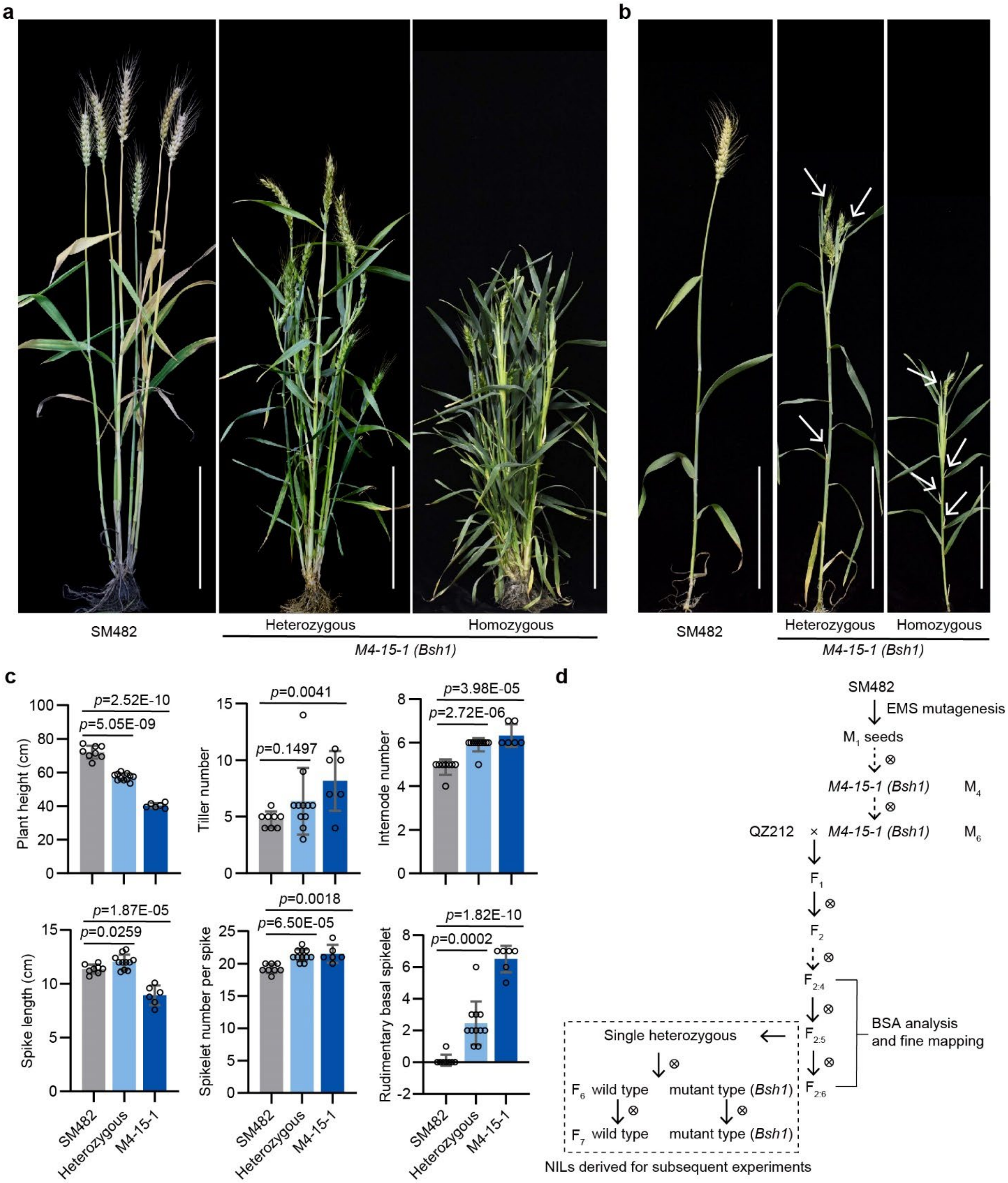


**Supplementary Fig. 1: Phenotypic characterization of the original *Bsh1* mutant line *M4-15-1* and the developmental workflow of genetic populations.**

**a**, **b**, Phenotypes of the wild type (SM482) and the original mutant line *M4-15-1* carrying the *Bsh1* allele. Plants segregating from the *M4-15-1* lineage exhibit distinct heterozygous and homozygous mutant architectures at the heading stage. **a**, Whole-plant architecture. **b**, Main culms showing ectopic axillary bud outgrowth and aerial branches. White arrows indicate ectopic aerial branches. Scale bars, 20 cm. **c**,

Comparison of agronomic traits between SM482 and the segregating *M4-15-1* mutants (heterozygous and homozygous), including plant height, tiller number, internode number, spike length, spikelet number per spike, and rudimentary basal spikelet number. Data are means ± s.d. (n ≥ 6 biologically independent plants). *P* values were determined by two-sided Student's *t*-test. **d**, Schematic overview of the origin of the *M4-15-1* mutant (via EMS mutagenesis of SM482) and the strategic workflow used to generate the mapping populations and near-isogenic lines (NILs). An initial $F_2$ population derived from a cross between *M4-15-1* and QZ212 was used to confirm the inheritance pattern. Subsequently, Bulked Segregant Analysis (BSA) was performed on extreme phenotypic pools explicitly selected from the advanced $F_4$ generation. For high-resolution fine mapping, advanced segregating families ($F_{2:4}$ to $F_{2:6}$) were utilized. The NILs carrying the *Bsh1* allele were subsequently derived from the selfed progeny of a single $F_{2:5}$ residually heterozygous individual and served as the primary genetic materials for subsequent physiological and molecular analyses.

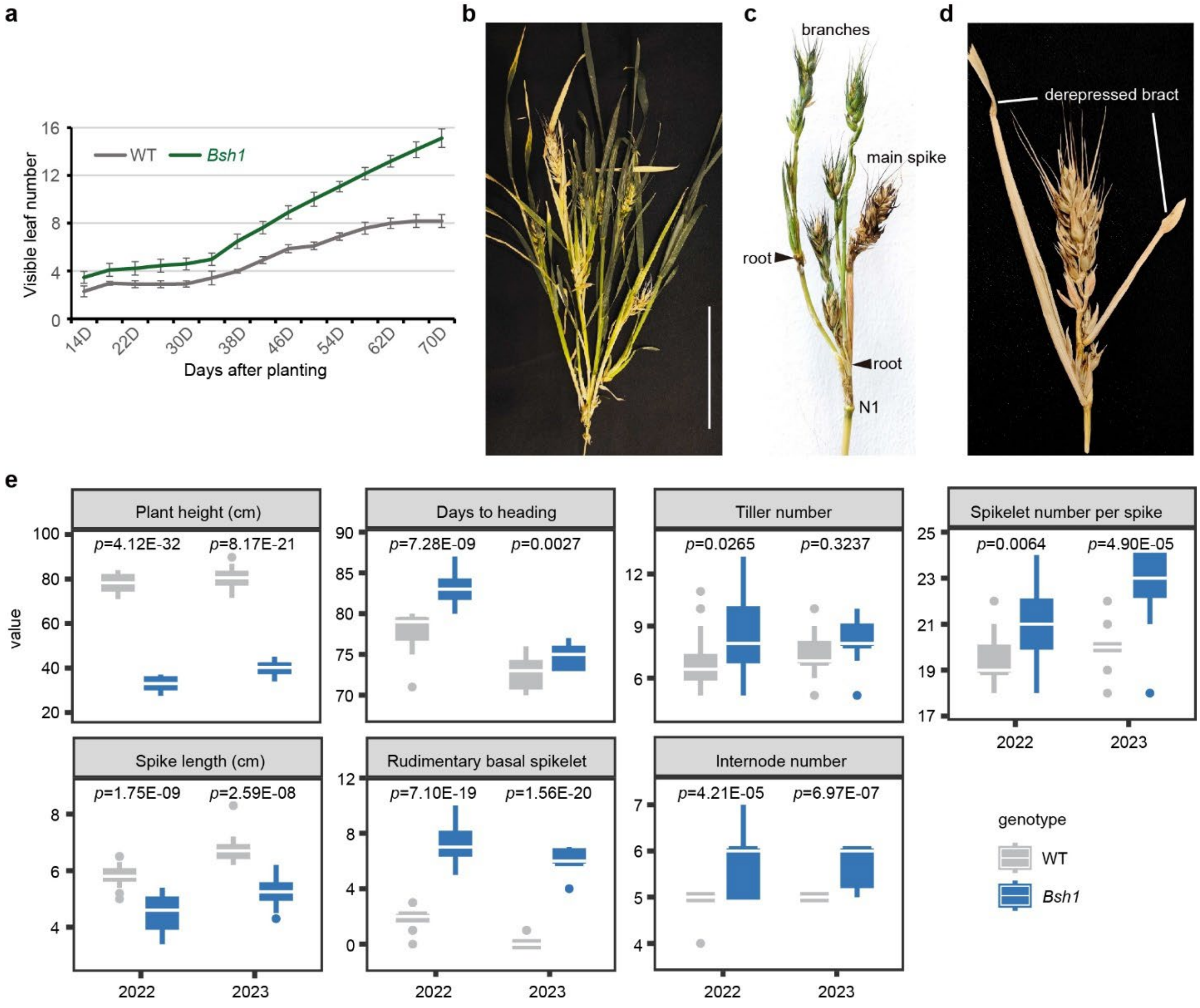


**Supplementary Fig. 2: Extended phenotypic and agronomic characterization of *Bsh1*.**

**a**, Number of visible leaves on the main shoot ($n \geq 40$ biologically independent plants per timepoint). **b–d**, Representative morphological defects in homozygous *Bsh1* plants. **b**, Mature shoot showing higher-order branches. Scale bar, 20 cm. **c**, Peduncle node showing ectopic aerial roots (black arrowheads) and axillary branches. **d**, Spike exhibiting derepressed bract growth (white lines) at the base of the rachis. **e**, Statistical comparison of agronomic traits, including plant height, days to heading, tiller number, spikelet number per spike, spike length, rudimentary basal spikelet number, and internode number ($n \geq 10$ biologically independent plants). In the box plots, the center line represents the median, box limits indicate the upper and lower quartiles, and whiskers extend to 1.5× the interquartile range. *P* values were determined by two-sided Student's *t*-test.

a b

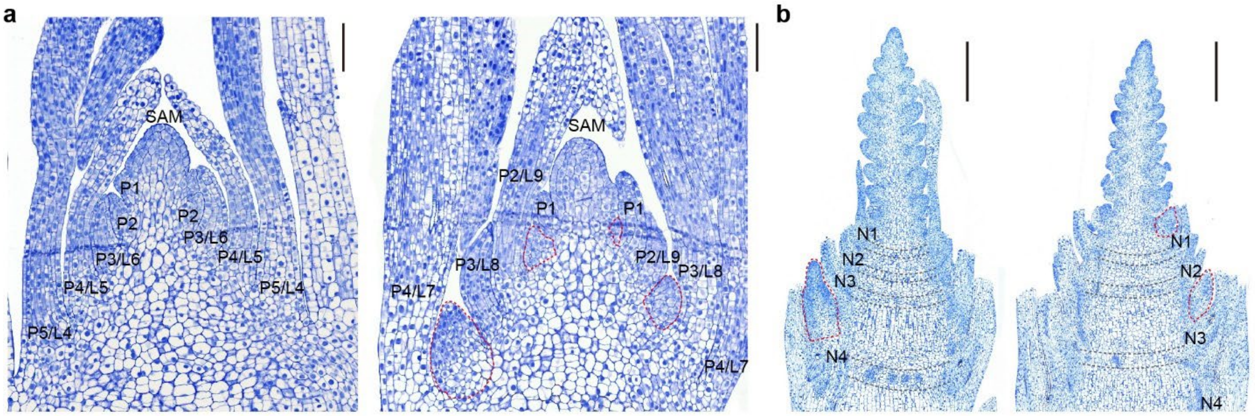


c

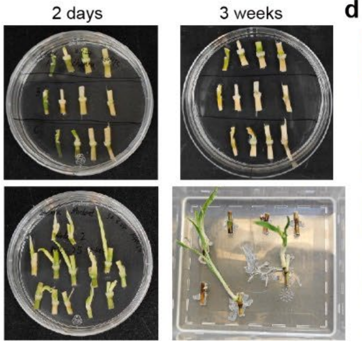


d

| | Sampling number | Contaminated sample | Transplanted number | Final plant number | Percentage (%) |
|---|---|---|---|---|---|
| *Bsh1* N0 | 30 | 5 | 25 | 0 | 0 |
| *Bsh1* N1 | 30 | 8 | 22 | 7 | 31.82 |
| *Bsh1* N2 | 30 | 11 | 19 | 1 | 5.26 |
| *Bsh1* N3 | 30 | 15 | 15 | 6 | 40.00 |
| WT N0 | 10 | 1 | 9 | 0 | 0 |
| WT N1 | 10 | 1 | 9 | 0 | 0 |
| WT N2 | 10 | 1 | 9 | 0 | 0 |
| WT N3 | 10 | 4 | 6 | 0 | 0 |

**Supplementary Fig. 3: Histological characterization of ectopic meristems and regenerative competency of upper culm nodes in *Bsh1*.**

**a**, **b**, Longitudinal sections of shoot apices from the wild type (WT; left) and *Bsh1* (right). **a**, Vegetative stage (12 days after germination). Red dashed outlines mark precocious axillary meristem initiation in *Bsh1* leaf axils, which is not detected in WT at this stage. **b**, Early reproductive stage (~W3). Ectopic axillary buds (red dashed outlines) are clearly visible and differentiating at the upper nodes (N1–N3) of *Bsh1*, whereas the corresponding regions in the WT remain smooth and dormant. SAM, shoot apical meristem; P, leaf primordium; N, node. Scale bars, 100 µm in (**a**) and 500 µm in (**b**). **c**, **d**, Regeneration assay to assess the developmental competence of upper culm nodes. **c**, Stem segments containing a single node were excised and cultured *in vitro*. Representative images show that after 3 weeks, *Bsh1* nodes (bottom) readily broke dormancy and produced shoots and roots, whereas WT nodes (top) remained quiescent or became necrotic. **d**, Regeneration efficiency for nodes N0-N3. N0 denotes the basal rachis node immediately subtending the spike; N1–N3 denote the successive nodes below. *Bsh1* upper nodes (N1–N3) exhibit strong regenerative competence, a trait largely absent in WT.

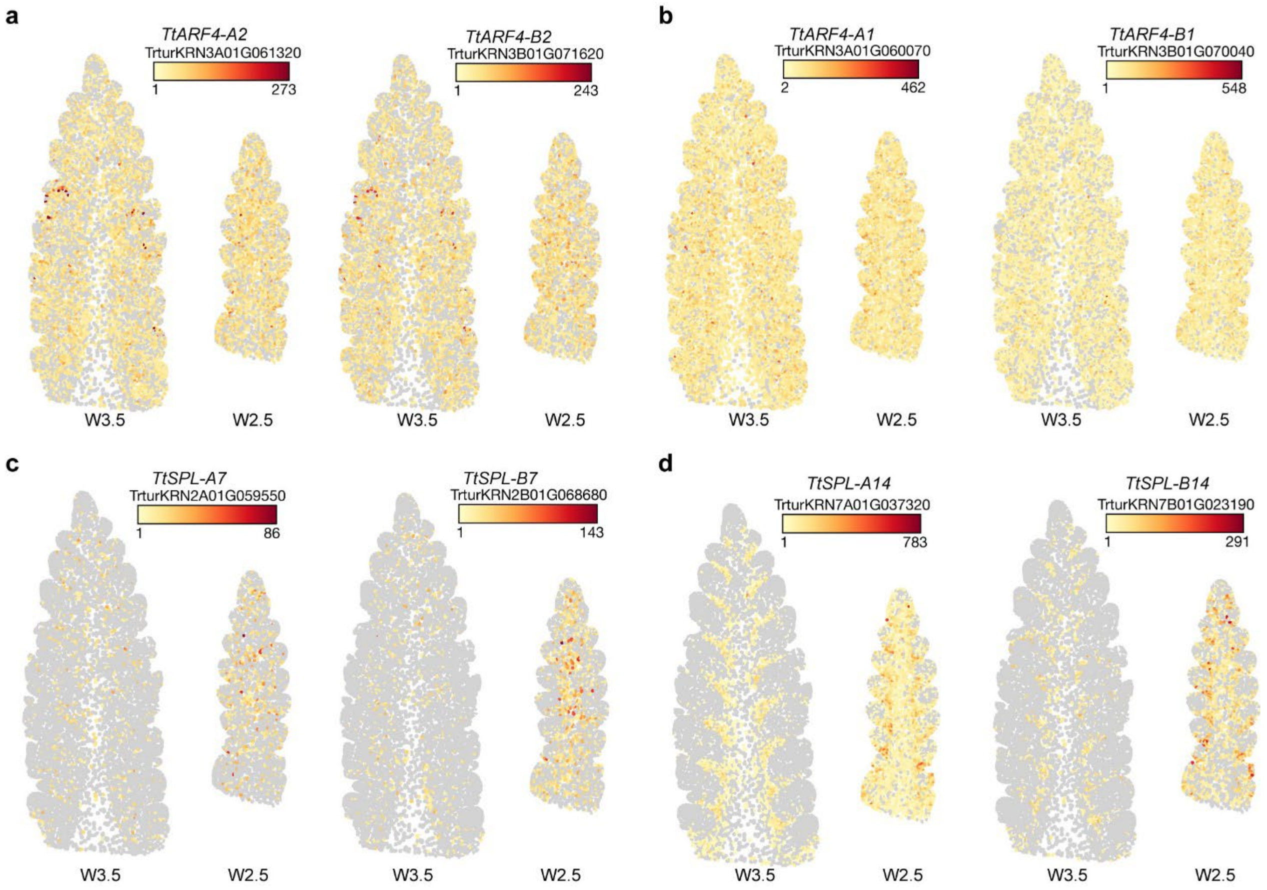


**Supplementary Fig. 4: Spatial expression landscapes of *ARF4* and *SPL* orthologs in tetraploid wheat spikes.**

**a–d,** Imputed spatial expression patterns of *ARF4* and *SPL* family genes in young spikes of tetraploid wheat (*Triticum turgidum* ssp. *durum* cv. Kronos) at the double ridge (W2.5) and floret primordium (W3.5) stages, derived from a published spatial transcriptomics dataset (Xu et al., 2025). **a,b,** Expression of *TaARF4-A2* orthologs *TtARF4-A2* and *TtARF4-B2* (**a**), and their paralogs *TtARF4-A1* and *TtARF4-B1* (**b**). These genes show broad expression across developing inflorescence meristems, consistent with *in situ* hybridization results in hexaploid wheat. **c,d,** Expression of *SPL* genes *TtSPL7* (**c**) and *TtSPL14* (**d**), showing overlap with ARF4 expression in bract tissues. *TaSPL14* (orthologous to *TtSPL14*) is identified as a direct downstream target of TaARF4-A2 in this study. Color bars indicate relative expression (imputed UMI counts; yellow, low; red, high). Trtur gene IDs are shown above each panel.

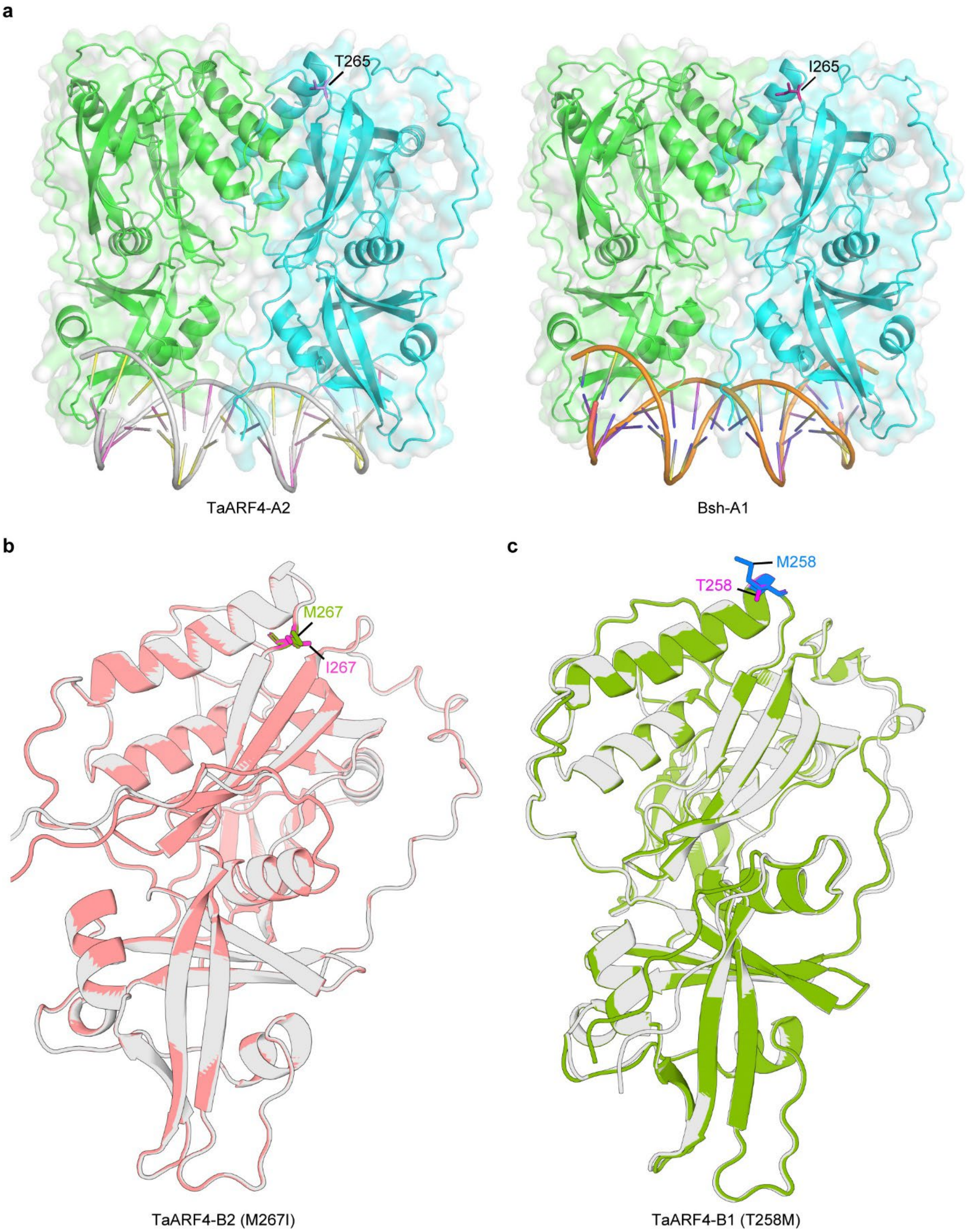


**Supplementary Fig. 5: Structural predictions of wild-type and mutant TaARF4 DNA-binding domains.**

**a,** Surface and ribbon representations of the predicted homodimer-DNA complexes for the wild-type TaARF4-A2 (left) and the Bsh-A1 mutant (right), generated using AlphaFold3 Server. Monomers are colored green and cyan. Residue 265 (T265 in WT, I265 in Bsh-A1) is indicated as sticks. This residue is located on an exposed loop distal to the DNA-binding interface. The overall predicted dimer architecture and DNA-interaction surfaces remain highly conserved, suggesting that the hypermorphic activity of Bsh-A1 is unlikely to arise from major global conformational changes in the DNA-binding domain (DBD). **b, c,** Superimposition of the predicted monomeric DBD structures for the wild-type TaARF4 homoeologs (grey) and their respective TILLING mutants. b, TaARF4-B2 and the adjacent-residue mutant M267I (pink). The M267 and I267 residues are highlighted as sticks. c, TaARF4-B1 and the adjacent-residue mutant T258M (green). The T258 and M258 residues are highlighted as sticks. Consistent with the T265I substitution, these structural alignments indicate that substitutions at these flanking conserved sites do not detectably alter the local secondary structure or overall protein folding.

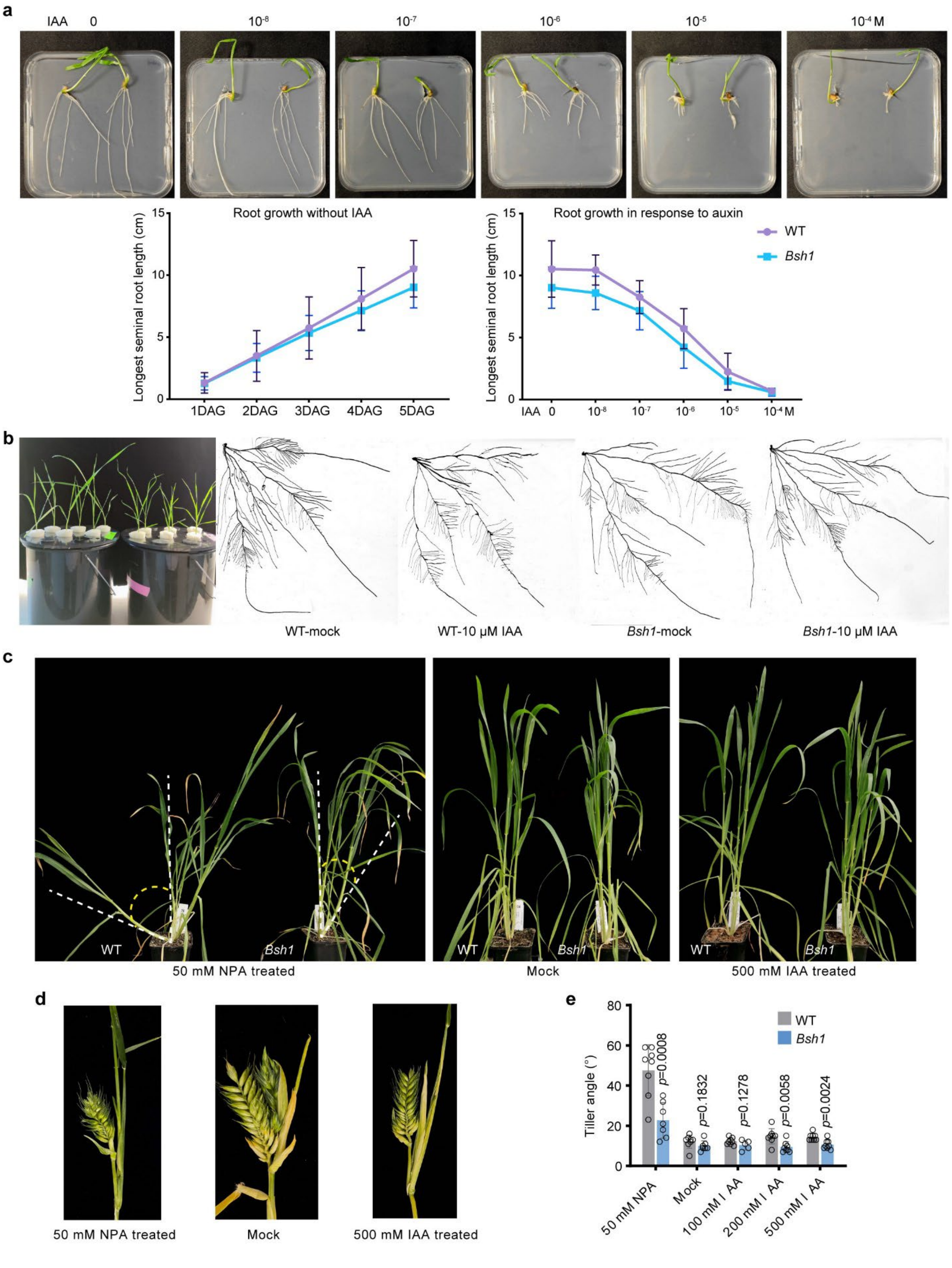


**Supplementary Fig. 6: Auxin sensitivity assays and pharmacological treatment experiments.**

**a,** Seminal root elongation assay on vertical agar plates. Top, representative seedlings grown on plates supplemented with indole-3-acetic acid (IAA; 0, $10^{-8}$, $10^{-7}$, $10^{-6}$, $10^{-5}$, $10^{-4}$ M). Bottom, seminal root length over time without IAA (left) and dose-response inhibition curves at 5 days after germination (DAG; right). Data are means ± s.d. (n = 5). Seminal root growth inhibition by IAA is comparable between WT and *Bsh1*. **b,** Root system architecture analysis in hydroponics. Representative whole plants (left) and scanned root systems (right) treated with mock solution or 10 mM IAA for 11 days. IAA strongly inhibits lateral root elongation in WT, whereas *Bsh1* lateral roots remain elongated. **c–e,** Pharmacological treatments with IAA or the auxin transport inhibitor N-1-naphthylphthalamic acid (NPA). **c,** Representative phenotypes after root drenching with Mock, 50 μM NPA, or 500 μM IAA. Dashed lines indicate tiller angle. Yellow arcs highlight increased tiller angle in NPA-treated WT, which is attenuated in *Bsh1*. **d,** Representative spikes from treated plants shown in **c**. Neither NPA nor high IAA rescue ectopic branching or bract outgrowth in *Bsh1*. **e,** Quantification of tiller angle under the indicated treatments. Data are means ± s.d. (n ≥ 5). *P* values were determined by two-sided Student's *t*-test.

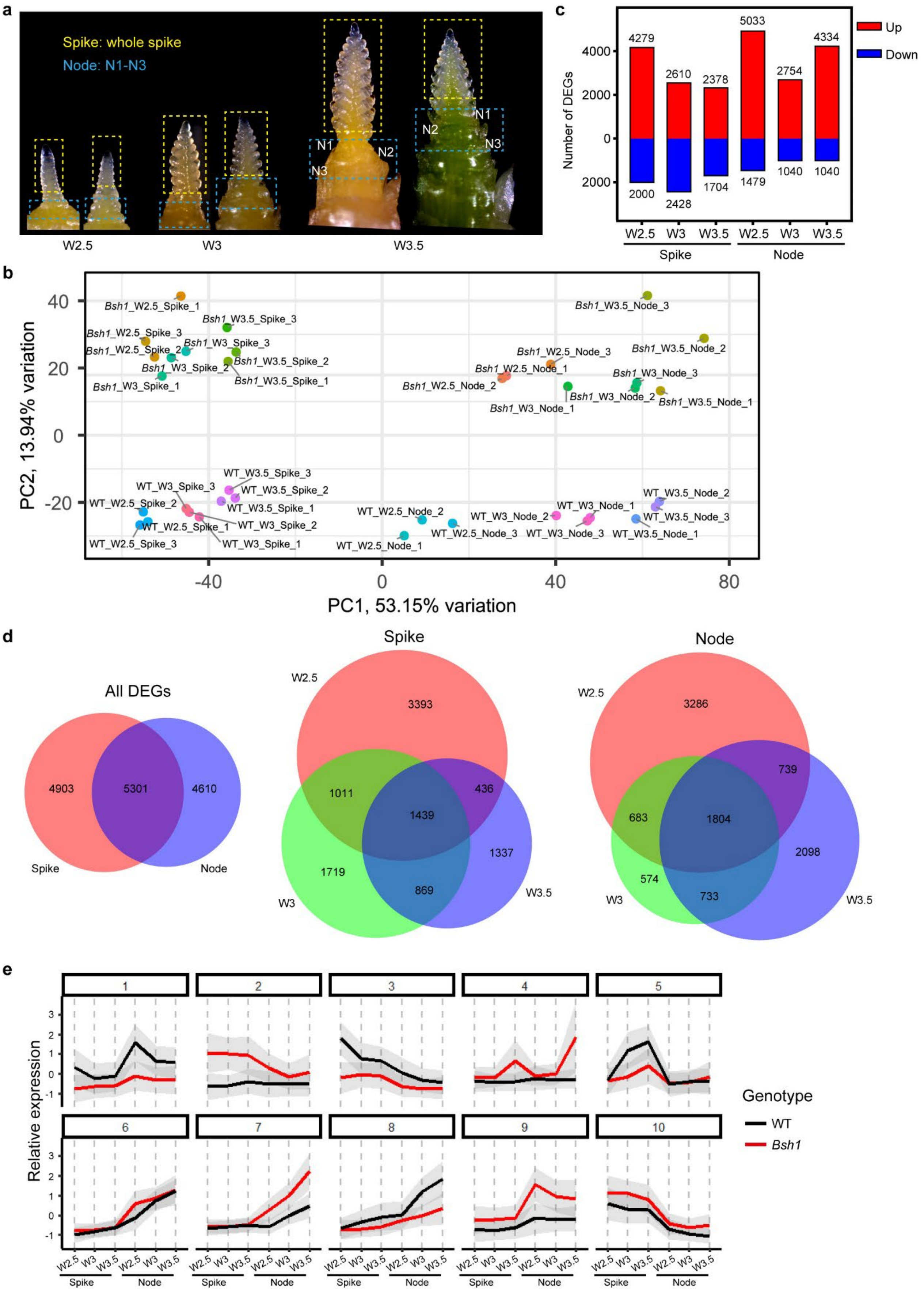
a
Spike: whole spike
Node: N1-N3
N1
N2
N3
W2.5
W3
W3.5
c
Number of DEGs
Up
Down
4279
2610
2378
5033
2754
4334
2000
2428
1704
1479
1040
1040
Spike
Node
b
PC1, 53.15% variation
PC2, 13.94% variation
Bsh1_W2.5_Spike_1
Bsh1_W2.5_Spike_2
Bsh1_W2.5_Spike_3
Bsh1_W3_Spike_1
Bsh1_W3_Spike_2
Bsh1_W3_Spike_3
Bsh1_W3.5_Spike_1
Bsh1_W3.5_Spike_2
Bsh1_W3.5_Spike_3
Bsh1_W2.5_Node_1
Bsh1_W2.5_Node_2
Bsh1_W2.5_Node_3
Bsh1_W3_Node_1
Bsh1_W3_Node_2
Bsh1_W3_Node_3
Bsh1_W3.5_Node_1
Bsh1_W3.5_Node_2
Bsh1_W3.5_Node_3
WT_W2.5_Spike_1
WT_W2.5_Spike_2
WT_W2.5_Spike_3
WT_W3_Spike_1
WT_W3_Spike_2
WT_W3_Spike_3
WT_W3.5_Spike_1
WT_W3.5_Spike_2
WT_W3.5_Spike_3
WT_W2.5_Node_1
WT_W2.5_Node_2
WT_W2.5_Node_3
WT_W3_Node_1
WT_W3_Node_2
WT_W3_Node_3
WT_W3.5_Node_1
WT_W3.5_Node_2
WT_W3.5_Node_3
d
All DEGs
Spike
Node
4903
5301
4610
3393
1011
1439
436
1719
869
1337
3286
739
683
1804
2098
574
733
e
Relative expression
Genotype
WT
Bsh1
Spike
Node

**Supplementary Fig. 7: RNA-seq experimental design and global transcriptome profiling.**

**a,** Sampling strategy for RNA-seq. Representative shoot apices at the W2.5 (double ridge), W3 (glume primordium), and W3.5 (floret primordium) stages. Dashed boxes indicate the precise tissues dissected for library construction: developing spikes (yellow) and subtending upper node tissues (N1–N3, blue) from WT and *Bsh1*. **b,** Principal component analysis (PCA) of RNA-seq datasets. PC1 and PC2 explain 53.15% and 13.94% of total variance, respectively. Samples cluster by genotype, tissue, and stage, indicating high reproducibility among biological replicates. **c,** Numbers of differentially expressed genes (DEGs) in *Bsh1* relative to WT across tissues and stages. Up-regulated (red) and down-regulated (blue) DEGs are indicated. **d,** Venn diagrams showing overlap of DEGs between spike and node tissues (left) and across developmental stages within spikes (middle) and node (right). **e,** K-means clustering of DEGs into 10 expression clusters across the six sample types. The x-axis indicates sample groups; the y-axis indicates Z-score-normalized expression. Black lines show WT centroids and red line show *Bsh1* centroids; grey shading indicates s.d.

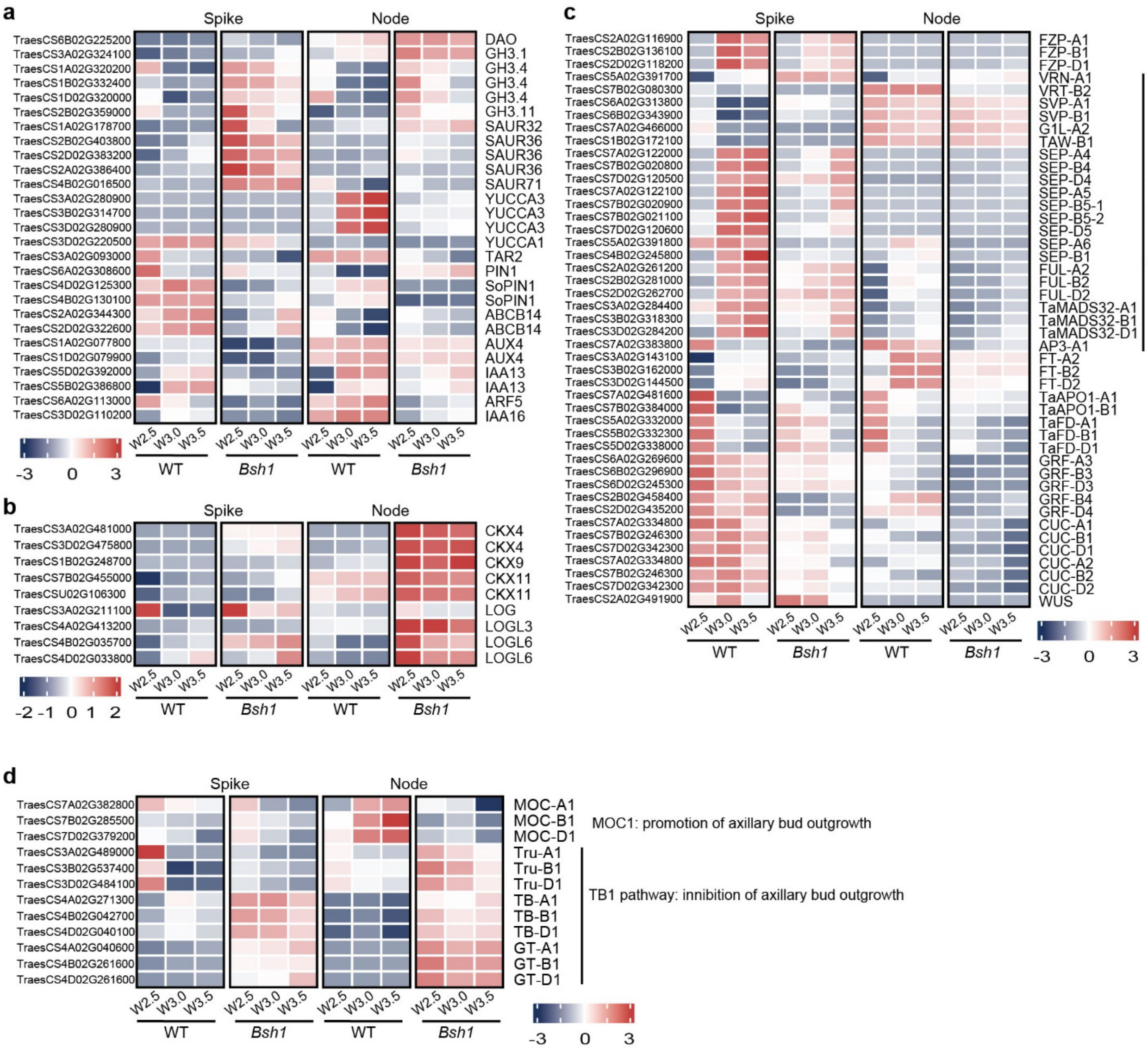


**Supplementary Fig. 8: Expression profiles of key DEGs involved in hormone signaling and developmental regulation.**

**a–d,** Heatmaps showing row-scaled expression (Z-scores) of selected DEGs in WT and *Bsh1* spikes and nodes at W2.5, W3.0, and W3.5. **a,** Auxin biosynthesis, transport, and signaling genes, including *YUCCA*/*TAR*, *PIN*/*SoPIN*/*ABCB*, and signaling response genes (*IAA*, *GH3*, *SAUR*, *ARF*). **b,** Cytokinin metabolism, including *CKX* and *LOG* family members. **c,** Inflorescence meristem and floral-development regulators, including MADS-box genes, WUSCHEL (*WUS)* and *CUP-SHAPED COTYLEDON* (*CUC*) family genes. **d,** Genes implicated in axillary bud outgrowth pathway, including *MOC1* and *TB1*/*GT1* modules. Gene IDs correspond to the Chinese Spring reference genome (IWGSC RefSeq v1.1).
